# Early memory deficits and extensive brain network disorganization in the *App*^*NL-F*^/*MAPT* double knock-in mouse model of familial Alzheimer’s disease

**DOI:** 10.1101/2021.04.11.439177

**Authors:** Christopher Borcuk, Céline Héraud, Karine Herbeaux, Margot Diringer, Élodie Panzer, Jil Scuto, S Hashimoto, T Saido, T Saito, Romain Goutagny, Demian Battaglia, Chantal Mathis

## Abstract

A critical challenge in current research on AD is to clarify the relationship between early neuropathology and network dysfunction associated to the emergence of subtle memory alterations which announce disease onset. In the present work, the new generation *App*^*NL-F*^/*MAPT* double knock in (dKI) model was used to evaluate early stages of AD. The initial step of tau pathology was restricted to the perirhinal-entorhinal region, sparing the hippocampus. This discrete neuropathological sign was associated with deficits in the object-place associative memory, one of the earliest recognition memories affected in individuals at risk for developing AD. Analyses of task-dependent c-Fos activation was carried out in 22 brain regions across the medial prefrontal cortex, claustrum, retrosplenial cortex, and medial temporal lobe. Initial hyperactivity was detected in the entorhinal cortex and the claustrum of dKI mice. The retention phase was associated to reduced network efficiency especially across cingulate cortical regions, which may be caused by a disruption of information flow through the retrosplenial cortex. Moreover, the relationship between network global efficiency and memory performance in the WT could predict memory loss in the dKI, further linking reduced network efficiency to memory dysfunction. Our results suggest that early perirhinal-entorhinal pathology is associated with local hyperactivity which spreads towards connected regions such as the claustrum, the medial prefrontal cortex and ultimately the key retrosplenial hub which is needed to relay information flow from frontal to temporal lobes. The similarity between our findings and those reported in the earliest stages of AD suggests that the *App*^*NL-F*^/*MAPT* dKI model has a high potential for generating key information on the initial stage of the disease.

## Introduction

Current diagnosis and treatment for Alzheimer‘s disease (AD) occurs too late, when physiological symptoms such as overt neurodegeneration have already reached an irreversible state (Selkoe, 2012). Hallmark pathologies of AD include insoluble amyloid plaques, initially formed in the medial prefrontal and posterior medial cortices indicative of the default mode network (DMN), and neurofibrillary tangles, initially formed in the medial temporal lobe (MTL). Combining functional neuroimaging with amyloid or tau PET reveals that spatial patterns of deposition induce functional network perturbations (Myers et al., 2014; Jones et al., 2017), and accompany cognitive decline (Sepulcre et al., 2017; Pereira et al., 2019). However, soluble amyloid peptide (Aβ) and hyperphosphorylated tau precursors are generally more toxic and accumulate in the brain decades prior to the onset of typical symptoms (Chen et al., 2017; Hill et al., 2020). This highlights the importance of focusing on the earliest stages to understand the origin of this devastating disease and to develop more proactive detection methods and therapies.

Amnestic mild cognitive impairment (aMCI) indicates a higher risk of progressing to AD and has been extensively studied over the last decade as a potential prodromal stage of AD (Petersen et al., 2009). Resting state functional connectivity (FC) has been evaluated in aMCI patients through the temporal correlation of signal fluctuations in fMRI, EEG, or MEG. In order to achieve a more rigorous understanding of FC disruption, graph theory has been used to evaluate the topology of functional networks. Healthy cognition likely requires efficient small world topology, combining local segregation between anatomically and functionally similar regions, and good integration between distant regions with the help of strongly connected hubs (Filippi et al., 2013, Bassett and Bullmore., 2006). In regards to aMCI, many studies have reported functional network perturbations within and between the DMN and the MTL, with reduced network integration along with reduced strength of cortical hubs (Drzezga et al., 2011; Wang et al., 2013a; Bai et al., 2013, Lin et al., 2019). However, these results are not entirely conclusive, as others have reported opposing or null results (Gardini et al., 2015; Grajski and Bressler, 2019; Liang et al., 2020, Liu et al., 2012). This inconsistency may be indicative of the heterogeneity of aMCI staging, where the sense of perturbation (hyperconnectivity vs hypoconnectivity) is shown to change depending on the severity of the pathological state (Pusil et al., 2019; Jones et al., 2016). This highlights the importance of identifying earlier biomarkers to more precisely detect preclinical stages of AD, even preceding aMCI. Recent progress has led to the development of sensitive recognition memory paradigms, such as pattern separation and associative memory tasks, which can detect slight cognitive defects in subjective cognitive impairment and cognitively healthy elderly (Naveh-Benjamin, 2000, Sinha et al., 2018; Reagh et al., 2018). These tasks depend on specific interactions across DMN and MTL structures sensitive to early AD (Reagh and Yassa., 2014; Miller et al., 2014; Hales and Brewer, 2011; Caviezel et al., 2020; Ritchey et al., 2020), and have be implemented to detect the earliest―asymptomatic‖ stages of preclinical AD (Rentz et al., 2011; Maass et al., 2019). By evaluating functional networks directly responsible for the earliest subtle memory deficits, we may achieve a clearer understanding of the neural networks first affected in the initial stages of AD.

Initial stages of AD are inevitably linked to discrete physiological and neuroanatomical perturbations, which should be more easily investigated in mouse models of AD (Scearce-Levie et al, 2020). Using fMRI on young AD transgenic mice, increased soluble Aβ (Shah et al, 2016; Shah et al., 2018), and regionally specific increases in neuroinflammation and phospho-tau (Degiorgis et al., 2020) have been linked to disruptions in resting state functional networks. As fMRI/EEG/MEG methods are currently difficult to implement in freely moving mice, ex-vivo imaging based on quantification of activity regulated expression of immediate early genes (IEG; c-fos, Egr1 and arc) may be used to evaluate memory related neural activation (Kinnavane et al, 2015). Using this technique, early-stage pathophysiology has been linked to individual changes in memory driven regional activity (Hamm et al., 2017), but not to functional networks. However, outside of AD, across-subjects correlations of activity regulated IEG levels have been used to assess functional networks directly related to memory (Tanimizu et al., 2017; Wheeler et al., 2013; Vetere et al., 2017). This approach can prove useful for evaluating memory driven FC in mouse models of AD.

Recently, knock-in mouse models have been created to express AD-related genes under endogenous mouse promotors (Saito et al., 2014). This removes gene overexpression and resulting potential artificial phenotypes, allowing for a more careful assessment of the initial stages of AD (Sasaguri et al., 2017). In the current study, we detected object-place (OP) associative memory deficits as the earliest sign of cognitive decline in the humanized *App*^*NL-F*^/*MAPT* double knock-in (dKI) mouse model. This model specifically expresses all six isoforms of tau and overexpresses Aβ_42_ which leads to marked amyloid deposition and typical pathological hyperphosphorylation of tau at an advanced age of 24 months (Saito et al., 2019). Given our specific interest in the early stages of AD, this model was chosen rather than the *App*^*NL-G-F*^*xMAPT* dKIs because of the slower development of its neuropathology. This option seemed fortunate as the first memory deficits detected in the OP task coincided with the emergence of the earliest stages of abnormal tau phosphorylation within the entorhinal-perirhinal region and not in the hippocampus, indicative of an early pathological state. Task-dependent activity was then measured in regions implicated with this form of associative memory, including primarily the DMN/MTL, and FC was assessed using graph theory techniques. Our results show that initial memory decline in AD could be linked to specific topological changes in memory dependent functional networks.

## Results

### Behavioural phenotyping reveals early alterations in object-place associative memory

Pattern separation (PS) and OP associative memory performances are commonly affected in prodromal stages of AD and cognitively normal elderly (Yeung et al., 2018, Hampstead et al., 2018, Reagh et al., 2018, Reagh and Yassa, 2014), and in mouse models at pre-plaque deposition stages (Zhu et al., 2017; Hamm et al., 2017). In order to detect earliest sensitive recognition memory deficits, a preliminary study was made to evaluate OP and PS performance, as well as long term object recognition performance at 2, 4, and 6 months of age (Figure 1—figure supplement 1). For OP and PS tasks, a short inter-trial-interval (ITI) of 5 minutes was chosen to ensure that deficits did not arise from a confounding perturbation of long-term memory consolidation. Given the high number of ages being tested and the difficult nature of the tasks, no results were significant in this preliminary experiment with multiple comparisons corrections. All groups succeeded in the long-term object recognition task and the PS task remained inconclusive because dKI‘s performances were highly variable (Figure 1—figure supplement 1A and 1B, respectively). On the other hand, a robust potential deficit could be seen at 4 months in the OP task, and persisted at 6 months of age (Figure 1—figure supplement 1C).

With a second independent cohort of mice, we confirmed that dKI mice were unable to detect a new OP association at 4 months of age (Figure 1A) (two sample t-test, WT vs. dKI: t(20) = 2.85, P = 0.0098; one sample t-test, against chance: dKI t(10)=0.3095, p = 0.763). On the other hand, 4-month-old mice did not show any deficits in short term novel object recognition (Figure 1B) (two sample t-test, WT vs. dKI: t(19) = .0934, P = 0.9265; one sample t-test, against chance: all ts>3.94, ps<0.003) and object location tasks (Figure 1C) (two sample t-test, WT vs. dKI: t(20) = 1.052, P = 0.9173; one sample t-test, against chance: all ts>3.03, ps<0.013). These results confirmed that the OP deficit was due to specific perturbations in associative memory rather than any separate impairments in the object recognition or spatial recognition.

**Figure 1.**
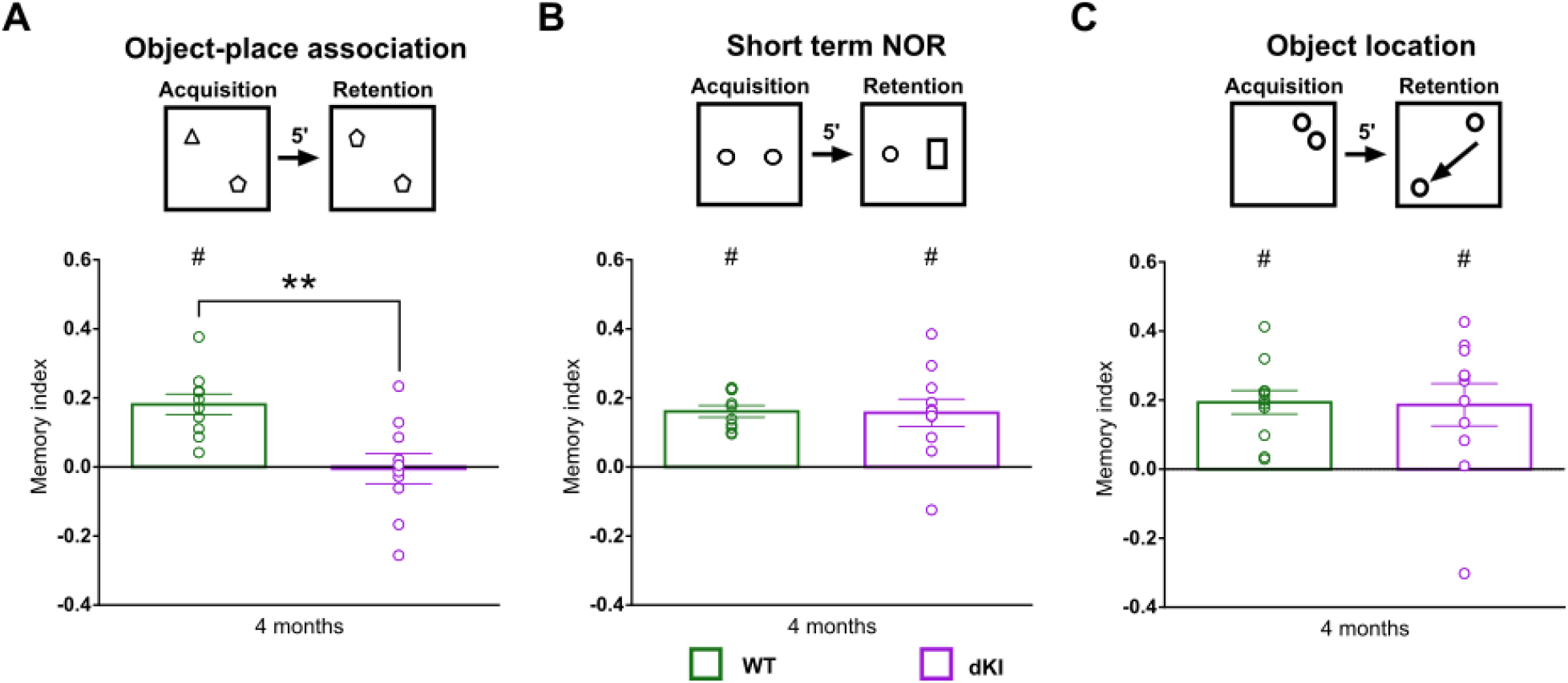
Behavioral characterization of young dKI mice (WT n = 11, dKI n = 11). (A) At 4 months of age, dKI mice reproduced the potential deficit seen in the object-place association task during preliminary phenotyping (two sample t-test, WT vs. dKI: t(20) = 2.85, P = 0.0098; one sample t-test, against chance: dKI t(10)=0.3095, p = 0.763). At the same age, dKI mice were normal in (B) object recognition (two sample t-test, WT vs. dKI: t(19) = .0934, P = 0.9265; one sample t-test, against chance: all ts>3.94, ps<0.003) (one WT mouse was removed due to being dropped before testing) and (C) spatial recognition (two sample t-test, WT vs. dKI: t(20) = 1.052, P = 0.9173; one sample t-test, against chance: all ts>3.03, ps<0.013), indicating that the loss in associative memory was not due to any singular loss in the short term object recognition or object location memory domains. Bar graphs represent the mean density (± SEM) * difference between genotypes **p<.01 (two sample t-test); #, difference from chance, #p<.05 (one sample t-test).

We then evaluated early AD pathophysiological changes in 4-month-old mice using western blotting (Figure 1—figure supplement 2A). Antibodies targeting the amyloid precursor protein, APP-cleaved carboxy terminal fragments, phosphorylated tau (Thr181) and total tau proteins were used. Wild-type mice expressed mainly APP α-CTF fragments whereas dKI expressed mainly APP β-CTF fragments. Moreover, an increase in β-CTF/α-CTF ratio was seen in both the hippocampus (Figure 1—figure supplement 2B) and the medial temporal cortex (MTC; including entorhinal, perirhinal, postrhinal cortices) (Figure 1—figure supplement 2C), indicating upregulated amyloidogenic APP processing. In addition, an increase of the tau phosphorylation degree on Thr181 was only seen in the MTC of dKI mice compared to wild-type mice while both models expressed similar levels of total tau proteins. This is consistent with general consensus of the MTC as the initial area of tau staging (Braak and Braak, 1995). These results highlight that OP deficits occur in conjunction with considerably light pathophysiological changes, akin to an early preclinical AD stage.

### Assessing OP dependent changes in brain activity

While encoding processes are known to be perturbed in AD patients (Granholm and Butters, 1988), recent studies in animal models also find dysfunctional retrieval of engrams as a mechanism for memory loss in early stages of AD-like pathology (Roy et al., 2016; Etter et al., 2019). We thus aimed to evaluate encoding versus recall dependent changes in brain activity in relation to the OP deficit, through independent evaluation of the two test-phases. Thus c-Fos activation was quantified in brains of mice that had underwent either the acquisition phase (*encoding*) (Figure 2A) or the retention phase (*recall*) (Figure 2B) of the OP task. In order to account for the c-Fos protein expression curve, the ITI was extended to 3 hours, the shortest delay that could permit definite isolation between test-phases. As with a 5-min ITI, dKI mice that underwent the retention phase after a 3-h ITI were likewise unable to detect the novel OP association (two sample t-test, WT vs. dKI: t(26) = 3.14, P = 0.004; one sample t-test, against chance: WT t(13) = 3.93, P = 0.002; dKI t(13) = 0.139, P = 0.891). (Figure 2C). We then chose to evaluate 22 regions of interest (ROIs) encompassing subregions of the medial prefrontal cortex (mPFC), claustrum (CLA), retrosplenial cortex (RSC), dorsal hippocampus (DH) and medial temporal cortex (MTC) (Figure 2D-J). Neuronal hyperactivity is common feature in early AD and in young pre-plaque mice (Zott et al., 2019). We therefore first assessed test-phase and genotype dependent changes in regional activity.

**Figure 2.**
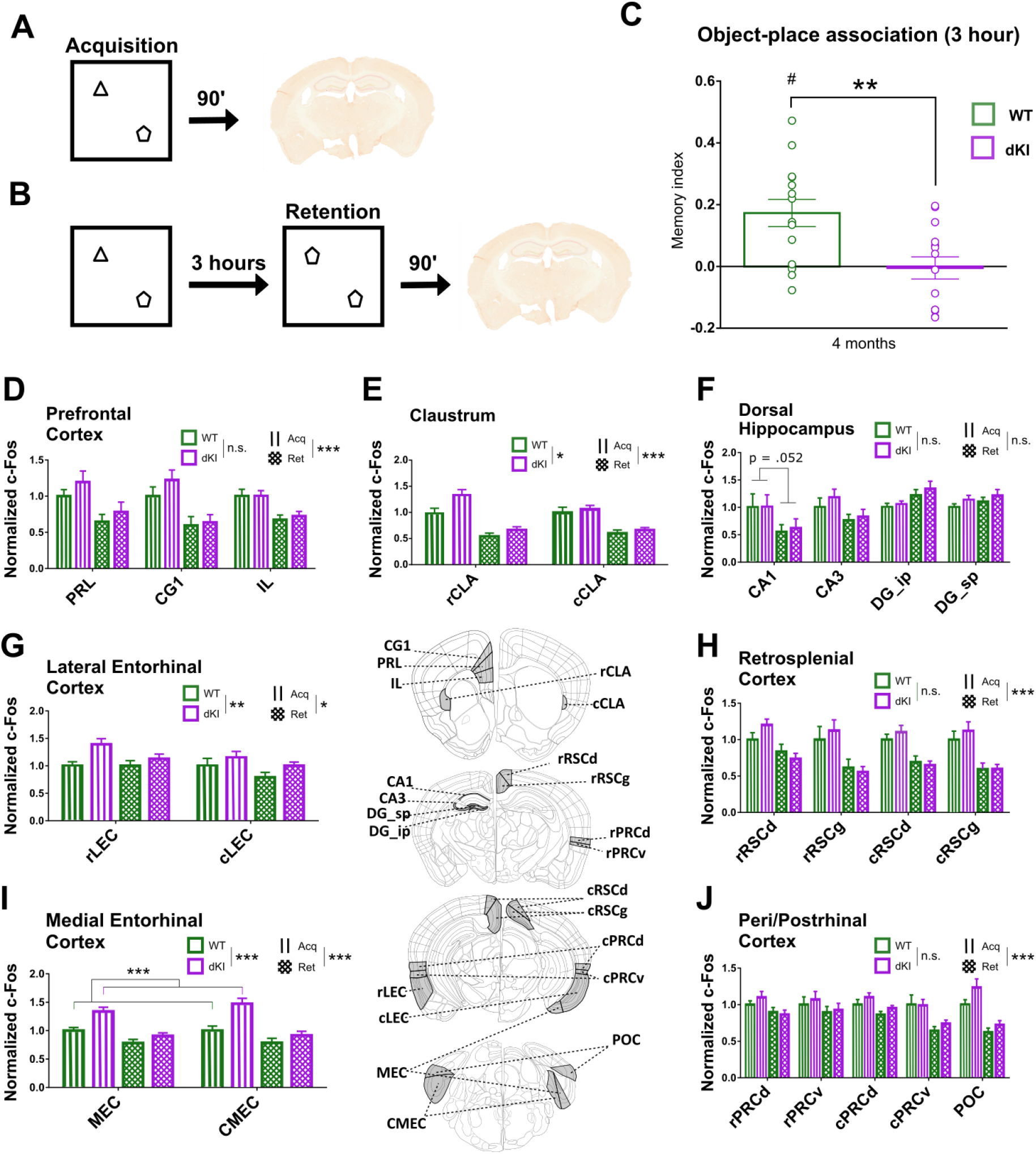
Regional activity during both phases of OP. Separate cohorts of WT and dKI mice were tested in either the (A) acquisition phase only or (B) the entire OP3h task, i.e., acquisition and retention phases. Ninety minutes following one test phase, brains were perfused and expression of the activity-regulated-gene, c-Fos, was evaluated immunohistochemically. (C) The dKI mice that underwent the whole OP3h task reproduced a deficit in object-place associative memory. difference between genotypes, **p<.01 (two sample t-test); difference from chance, #p<.05 (one sample t-test). (D-J) Graphs that illustrate c-Fos counts normalized to the WT-Acquisition group. For each region group a 3-factor ANOVA was performed for test-phase (Acquisition-Lines, Retention-Grid), genotype (WT-green, dKI-purple) and subregion effects. Significance for test-phase and genotype effects are depicted above each region graph; *p<.05, **p<.01, ***p<.001. (D-E,G-J) There was an increase in activity during the acquisition phase across all regions, except the (F) dorsal hippocampus. The (E) claustrum and (G) lateral entorhinal cortex presented increases in activity in dKI mice regardless of the test-phase. The (I) medial entorhinal cortex however, was hyperactive specifically during the acquisition phase ***p<.001 (Tukey post hoc). Bar graphs represent the mean density (± SEM). Subregion prefixes – r,rostral;c,caudal; suffixes – RSCd,dysgranular; RSCg, granular; PRCd, dorsal; PRCv, ventral; ip, infrapyramidal; sp, suprapyramidal. Error bars indicate the SEM. Schematics indicating the general locations of brain regions were adapted from Allen Mouse Brain Atlas derived vector images (Lein et al., 2007).

### Acquisition induces higher regional activity than retention

All regions except the DH (phase effect: DH F(1, 192) = 1.85, P = 0.175; subregion x phase effect: F(3, 192) = 5.32, p = 0.002; post-hoc Tukey: Acq vs Ret: CA1 P = .052, CA3 P = .383, DG_ip P = .597, DG_sp P = .513) showed higher levels of activity in the acquisition phase, although in the LEC this increase in activity was noticeably less drastic (phase effect: LEC F(1, 96) = 5.22, P = 0.025; all other Fs > 25.6, Ps < 0.001). This can be expected in the initial phase as the mice are subjected to stronger changes with two new objects in previously unoccupied locations in respect to the retention phase, where the only change is a new OP association. During acquisition, the mice are actively encoding several new features, new objects and their unique location in the open-field. Moreover, this helps confirm the absence of residual c-Fos in the retention groups. If residual c-Fos lingered from encoding until perfusion after the retention phase, one would expect similar or increased c-Fos density during the retention phase, not a decrease.

### Hyperactivity in the claustrum and entorhinal cortex of dKI mice

The CLA and LEC showed increased activation in dKI mice regardless of the test phase (genotype effect: CLA F(1, 96) = 8.29, P = 0.005; LEC F(1, 96) = 10.36, P = 0.001). The MEC revealed increased activation in dKI mice specifically during the acquisition phase (MEC: genotype effect: F(1, 96) = 25.64, P < 0.001; genotype x phase effect: F(1, 96) = 7.46, P = 0.007; post-hoc Tukey: WT vs dKI during acquisition P < .001, WT vs dKI during retention P = 0.252).

**Table 1.**
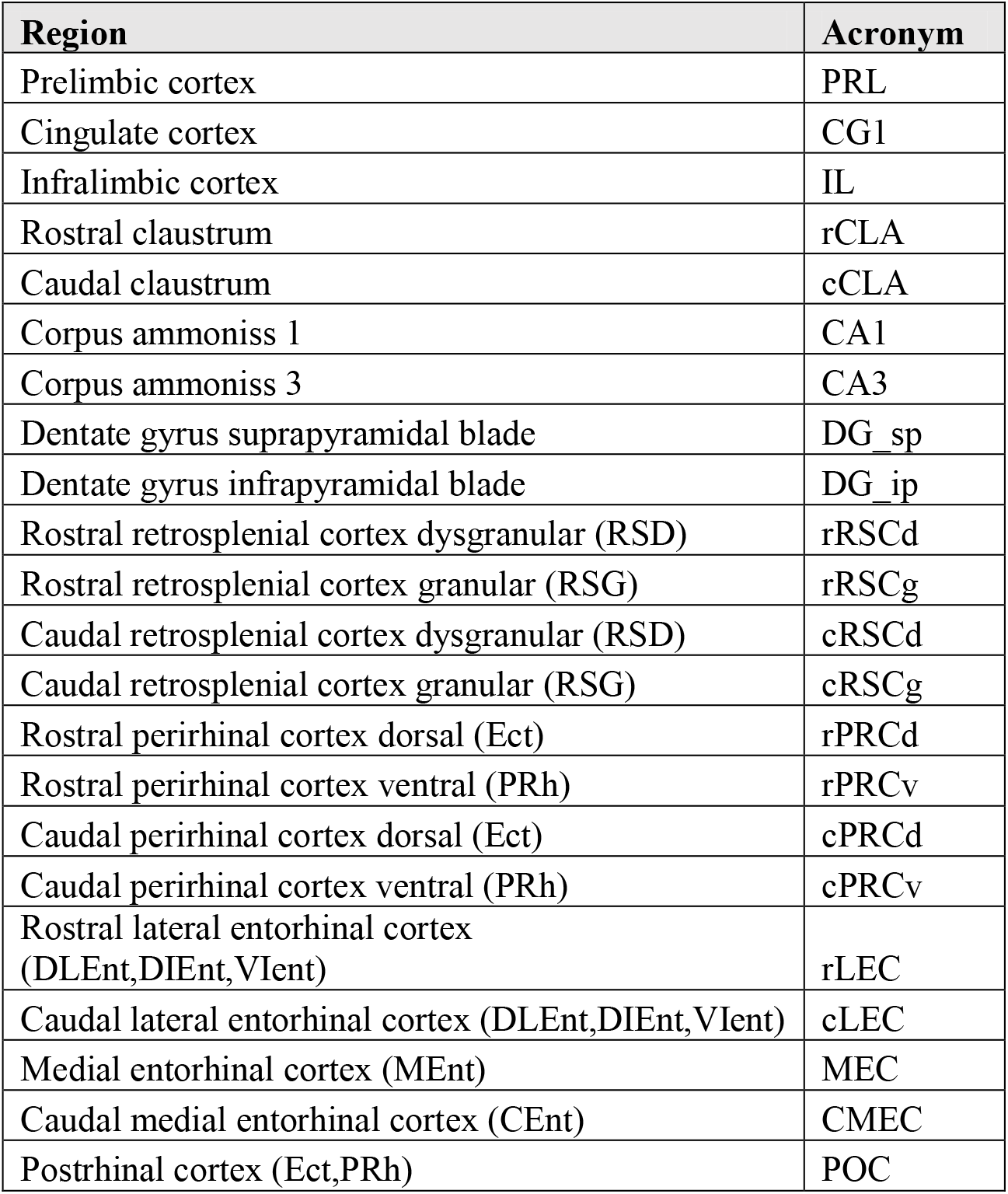
List of evaluated regions. Regions were chosen a priori based on their relevance to early AD pathology and to associative memory processing. In parentheses are region identifications according to the third edition Franklin and Paxinos Mouse Brain Atlas.

### Computing functional couplings

Brain regions with correlated activity can be said to exhibit functional connectivity (FC), as co-modulations of activity can be a marker of inter-regional information sharing. In humans or head-restrained/anesthetized rodents recorded with EEG, MEG (for electrical signals) or fMRI (for metabolic rate), FC is computed from the covariance across time. In methods using regional expression of c-Fos, FC can be modeled by the covariance of regional activity across subjects to study memory driven networks in mice (Wheeler et al., 2013; Tanimizu et al., 2017). The validity of this approach has been confirmed using chemo-genetic techniques (Vetere et al., 2017).

From the c-Fos signal, we assessed FC by computing the inter-regional Spearman correlation coefficients (r) for each Genotype-Phase group. Correlation matrices were used to visualize all possible correlations within each group (Figure 3A-B). We first assessed global FC strength by taking the mean r value of each matrix. Most FC couplings were positive, however we found a few weakly anti-correlated pairs of regions. In evaluating FC, there were 3 ways to consider these negative correlations; as disruptive, as very-weak or as contributing equally to positive correlations. All three cases were evaluated by taking the mean r with retained, near-zeroed, and absolute valued negative correlations respectively. In the near-zeroed case, negative correlations were reduced to a value of .006, the smallest positive correlation observed.

**Figure 3.**
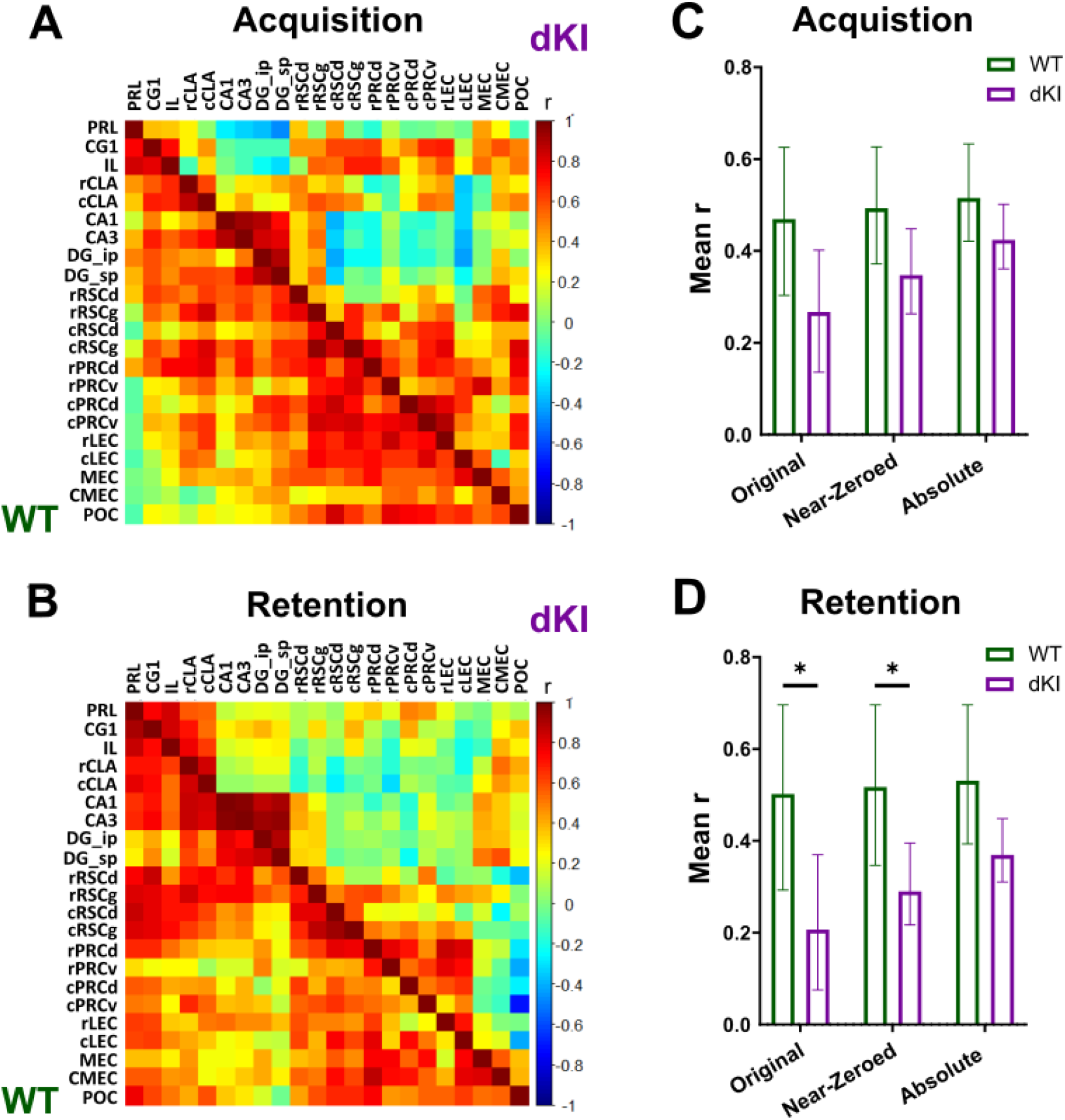
Functional connectivity is depicted with matrices showing across subjects inter-regional spearman correlations for c-Fos expression during (A) retention and (B) acquisition. Colors reflect correlation strength (scale, right). We assessed global FC strength by taking the mean r value of each matrix with retained, near-zeroed, and absolute valued negative correlations. In the near-zeroed case, negative correlations were reduced to a value of .006, the smallest positive correlation observed. (C) We found no change in global FC strength for any of the three cases during acquisition. (D) During retention however, there was a significant decrease in global FC strength with the original and near-zeroed correlations, but not when they are absolute valued. Bar graphs represent the mean bootstrap value, and the error bars represent the bootstrapped 95% confidence interval. * -the 95% CI for the difference ≥ 0

During acquisition, we found no significant change in global FC strength between WT and dKI groups, for any of the three ways to treat negative correlations in FC (Figure 3C). However, during retention, there was a decrease in global FC strength in dKI mice with respect to WT. This decrease could be proved significant when considering negative correlations as disruptive or as very weak, but not when their absolute value was taken (Figure 3D). This indicates that two phenomena coexist: first, a reduction of positive inter-regional correlations, corresponding to decreased ―cooperation‖ between some regions; second, an increase in absolute strength of inter-regional negative correlations, corresponding to increased―conflict‖ between some regions.

### Generating functional networks

Each functional connectivity matrix can be considered as the adjacency matrix of a weighted undirected network (Rubinov and Sporns, 2010), and, as such, its organization can be assessed through the analysis of functional networks, using techniques arising from graph theory. From each matrix a functional network was generated as a fully connected weighted graph. In these graphs, the regions are represented as nodes and the functional connections between regions are represented by edges. The strength of a connection between two regions, as represented by the edge weight, is determined by their inter-regional correlation strength. As negative correlations are weak and pose difficulties of interpretation when dealing with many graph theory techniques, we decided for all following analyses to replace them with near-zeroed values when constructing our functional networks, in line with other studies of c-Fos derived functional networks in-vivo (Vetere et al., 2017). It has been shown that community organization of structural networks can change between healthy individuals and different stages of MCI (Pereira et al., 2016). We decided to evaluate community organization of each functional network to see if any consistent or phase specific changes could be detected between genotypes. This could also aid us in obtaining a qualitative understanding of organization of information flow in each network. Network communities were detected through a modularity maximization procedure (see Materials and Methods). This unsupervised algorithm separates the nodes into distinct groups (the communities), such that nodes within a community are more strongly interacting between them than with nodes in other communities. The separation in communities will partly stochastically vary for different runs of the algorithm and different bootstrap instances of the FC matrix. To extract a robust consensus set of communities we then computed allegiance matrices (Bassett et al., 2015), which depict the percentage of times that any given pair of regions is attributed to the same community among bootstraps (Figure 4).

**Figure 4.**
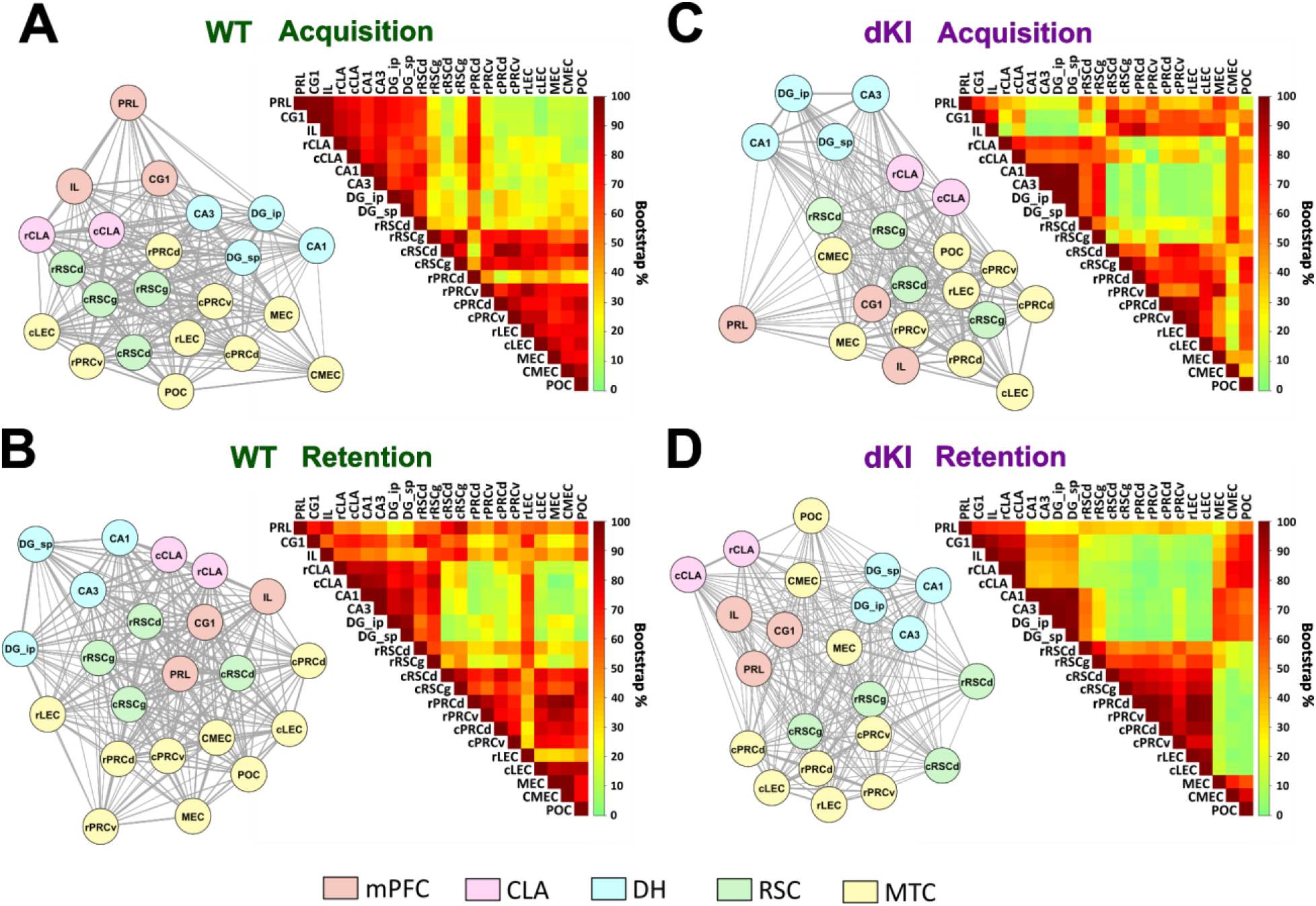
Funtional network structure and community organization. Functional networks were computed as fully connected weighted graphs, with regions as nodes (circles) and inter-regional correlation strengths as edge weights (line thickness). For visual examination of networks, nodes were placed so that they lie closer to nodes with which they are strongly connected, and were color coded according to a priori region subfields. Communities were detected through modularity maximization, which finds communities of regions that have stronger connections with each other relative to the rest of the network. Communities were detected across bootstrapped networks, and allegiance matrices were utilized to depict community stability as the percentage of bootstraps that contain any given pair of regions in the same community. (A,B) In both WT groups there is a stable MTC/cRSC community (red area at the bottom right of each WT allegiance matrix). (A) In the WT-Acq there is a stable DH/mPFC community (red area at the top left of the allegiance matrix). (B) In the WT-Ret the mPFC shares community allegiance with all of the network (red area along the mPFC rows across the top of the allegiance matrix) as mediated through the RSC (darker red square in the middle of the top three mPFC rows). (C,D) Across both dKI groups, the MEC/CMEC/POC regions appear to display a consistently modified community allegiance in respect to the WT, associating more with the DH/mPFC and sometimes less with the rest of the MTC (red area at top right, some green areas at bottom right of each dKI allegiance matrix).

### WT networks differentially engage the DH and mPFC

A stable community is found in both task phases encompassing most of the MTC and the cRSC (Figure 4A,B; red area at the bottom right of each WT allegiance matrix). The main difference between the WT-Acquisition and WT-Retention groups was the differential involvement of the DH or mPFC with this MTC community. During acquisition, the DH shared a community primarily with the mPFC (Figure 4A; red area at top left the allegiance matrix). During retention, the mPFC shared allegiance across the network (Figure 4B; red area along the mPFC rows across the top of the allegiance matrix). This more global incorporation of the mPFC appeared to be driven largely through the RSC (Figure 4B; darker red square area located at the middle of the top three mPFC rows).

### dKI networks reveal consistent departures from WT community structure

During acquisition, the DH community largely disengaged with the mPFC as compared to the WT-acquisition group (Figure 4C; green square at the top left of the allegiance matrix). Moreover, the DH appeared to be more strongly aligned to the CMEC, MEC, and POC as compared to the rest of the MTC (Figure 4C; the more red/orange area at the right of the DH rows).

During retention, both the mPFC and the DH disengaged with the MTC. This led to distinct communities containing mPFC/CLA, DH and the MTC (Figure 4D; three separate red squares of the allegiance matrix). Similarly to the acquisition phase, the MEC/CMEC/POC regions were heavily recruited by the mPFC and DH communities (Figure 4D; red area at top right), at the cost of disengagement with the MTC community (Figure 4D; green area at bottom right of the allegiance matrix). The consistent recruitment of the MEC/CMEC/POC in both dKI groups elucidates a possible role for these regions as compensatory hubs in this mouse model.

### Information flow in functional networks

Efficient integration of information flow across functional networks is shown to aid in cognitive function (Wang et al., 2013a; Martinez et al., 2018). Interpreting network links as ―pipes‖, how well can a network allow information to flow will depend not only on how wide individual pipes are, but also on how pipes are disposed and aligned to form pipelines between the nodes that must communicate, without too many steps and bottlenecks. The general capacity for a network to sustain efficient flows is quantified in graph theory by metrics such as *global efficiency* (see Materials and Methods; Latora and Marchiori, 2001). It is important to note that even if a network has reduced global FC strength, its global efficiency may rest unaffected if there are well placed ―hub‖ regions to facilitate indirect communication. To evaluate information flow at the regional level, *nodal strength* and *nodal efficienc*y metrics are used. Nodal strength measures the degree to which each specific region can exchange information directly with all other regions of the network. A region with low strength may still be able to communicate with its network indirectly, again likely through hub regions. This indirect communication can be measured using nodal efficiency. Organization of information flow in a network can be examined through strength and efficiency distributions. Regions with higher strength and efficiency can be considered to contribute more to their network. High direct connectivity of a region, such as high strength, is also shown to describe potential hub regions (Vetere et al., 2017) which, as we have described, are needed to facilitate indirect information flow and maintain global efficiency. We compared information flow between WT and dKI networks within each test-phase. Organization of information flow was first evaluated through a qualitative examination of node strength and nodal efficiency distributions. To best visualize distributions, the strength/efficiency values were squared and normalized to the highest value within each network, and were displayed as necklace diagrams (acquisition - Figure 5A,B) (retention – Figure 5F,G). We then directly compared global efficiency and nodal strength/efficiency between genotypes in subsequent analyses (acquisition - Figure 5C,D,E) (retention - Figure 5H,I,J).

**Figure 5.**
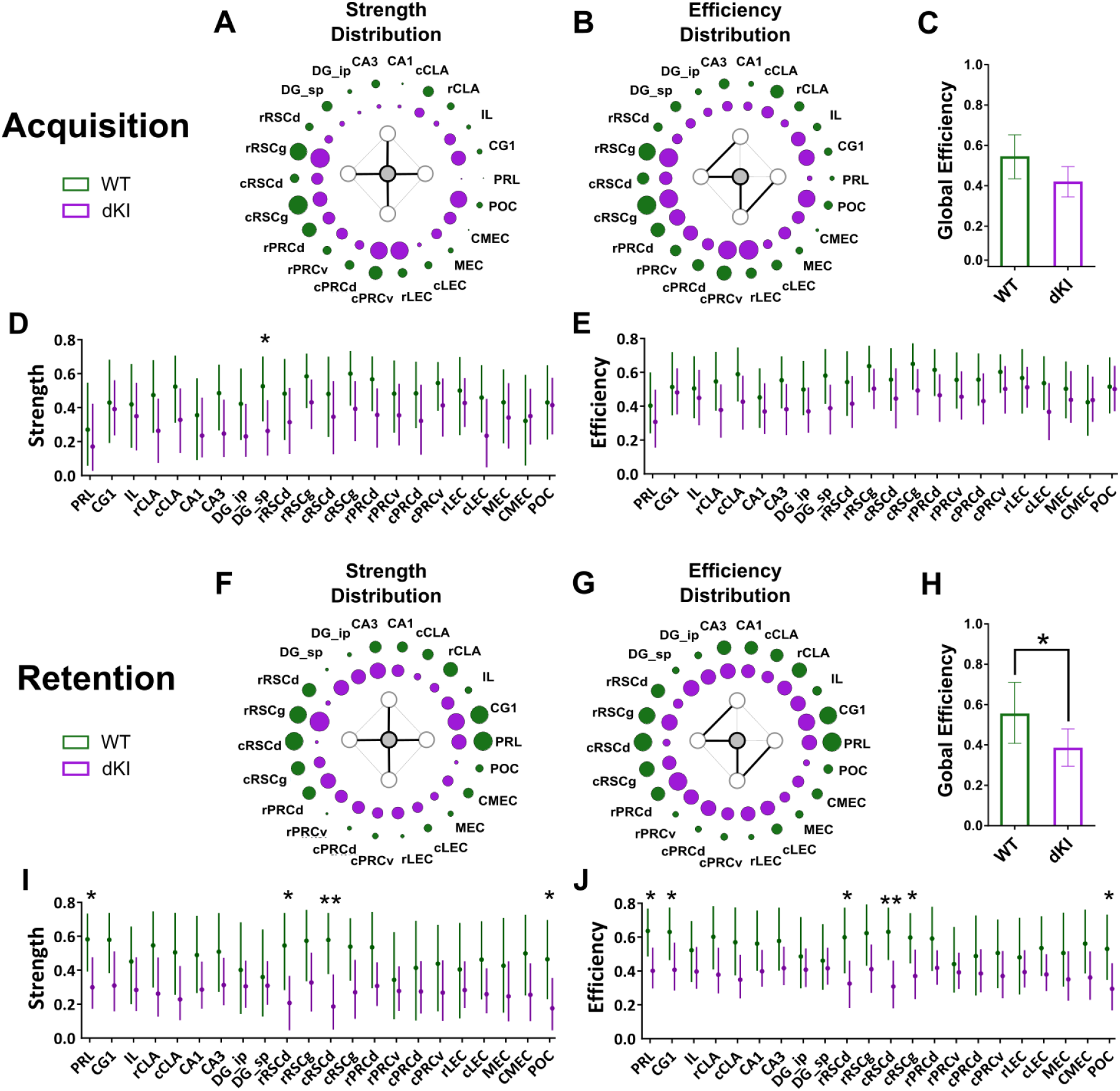
Information flow in functional networks. Network organization of information flow was assessed through examination of node strength and nodal efficiency distributions in both test-phases (acquisiton,retention). These distributions were visualized using necklace diagrams, where circle size reflects the within network normalized (A,G) strength^2^ and (B,H) efficiency^2^. Global efficiency was then compared directly between genotypes to assess (C,F) network integration. (D,I) Nodal strength and (E,J) nodal efficiency were compared between genotypes to assess region dependent changes in direct and indirect information flow, respectively. (A,B) During acquisition, strength and efficiency distributions were roughly conserved between genotypes, with strong emphasis on the RSC and MTC regions, although in the dKI, mPFC, MEC and POC subregions were much more involved. (C) Global efficiency was not severely affected in dKI mice. (D) A drop in region strength was seen in the DGsp, (E) but its efficiency was unaffected. (G,H) During retention, heavy emphasis was placed on mPFC and RSC regions in the WT network. In the dKI, this emphasis is largely lost. (F) Global efficiency of the dKI network was significantly reduced as compared to the WT network. (I) Reductions in region strength were seen in the POC, PRL, and most severely in both subregions of the RSCd. (J) Reductions in efficiency was retained among these regions, with additional losses in the CG1 and the cRSCg. Bar graphs and dot graphs represent the mean bootstrap value, and the error bars represent the bootstrapped 95% confidence interval. * -the 95% CI for the difference ≥ 0 ; ** -the 99% CI for the difference ≥ 0

### Acquisition– the DG_sp maintains efficiency despite a drop in strength

During acquisition, strength and efficiency distributions were largely conserved between genotypes with strong emphasis on RSC and MTC regions. Although, in the dKI there appeared to be additional involvement of mPFC, MEC and POC (Figure 5A,B). Nevertheless, global efficiency was not severely affected in the dKI network (Figure 5C). The unique significant drop in nodal strength was seen in the DG_sp (Figure 5D), but as there was no drop in nodal efficiency (Figure 5E), it could still effectively communicate with the rest of the network indirectly. This maintenance of indirect communication, despite drops in direct communication, reflects the potential importance of the rRSC or MEC/CMEC/POC compensatory hubs.

### Retention – severe losses in network efficiency, especially across the cingulate cortex

During retention, heavy emphasis was placed on mPFC and RSC regions in the WT network. In the dKI network this emphasis was largely lost and the distributions take on a more homogenous structure, though the rRSCg and mPFC subregions still appeared to be among the most involved (Figure 5F,G). Global efficiency was significantly reduced in the dKI network (Figure 5H), with sharp drops in strength seen in the PRL and the POC, but most severely in both subregions of the RSCd (Figure 5I). Loss in nodal efficiency was even more prevalent, with additional reductions in the CG1 and the cRSCg (Figure 5J). These results suggest that retention dependent functional integration across the cingulate cortex is severely disrupted in the dKI, and may be linked to reduced hub strength of the RSCd.

### Relating network efficiency and memory

Finally, we addressed the question of whether the reduction of OP memory performance in dKI mice could be accounted for by the reduction in network efficiency observed during retention. Our aim was thus to determine if a relationship could be found between global efficiency and memory index in the WT and be used to predict dKI memory loss. This was assessed through correlation testing across sub-sampled WT populations, where each sub-sample corresponded to the removal of one mouse. From each sub-sample the average memory index and resulting global efficiency were calculated, and correlation significance was tested with Pearson and Spearman correlation coefficients.

### Global efficiency positively correlates with an exploration adjusted memory index, and predicts memory deficiency in the dKI

No significant correlation was found between global efficiency and memory index across WT sub-samples (Figure 6 - figure supplement 1A). However, we noticed that of the mice with high memory index, those with greater exploration times (total time exploring both objects) appeared to have a weaker contribution to global efficiency. This hinted a potential dampening effect of general object exploration on memory driven FC. Note that increases in exploration time themselves may be interpreted as a further sign of memory function alteration, beyond reductions in the previously defined memory index MI. We thus defined an exploration index EI, quantifying the exploration time of the mouse normalized to the group and used it to compute as well an exploration adjusted memory index as the memory index divided by the exploration index (MI/EI). Decrease of such an exploration adjusted memory index may reflect both MI decrease and EI increase, thus summarizing both probed facets of memory-related behavior alteration. As in the case of MI alone, we did not find a significant correlation between global efficiency and EI index (Figure 6 - figure supplement 1B). A significant positive correlation was found however between global efficiency and the adjusted memory index MI/EI across the WT sub-sampled networks (Figure 6).

**Figure 6.**
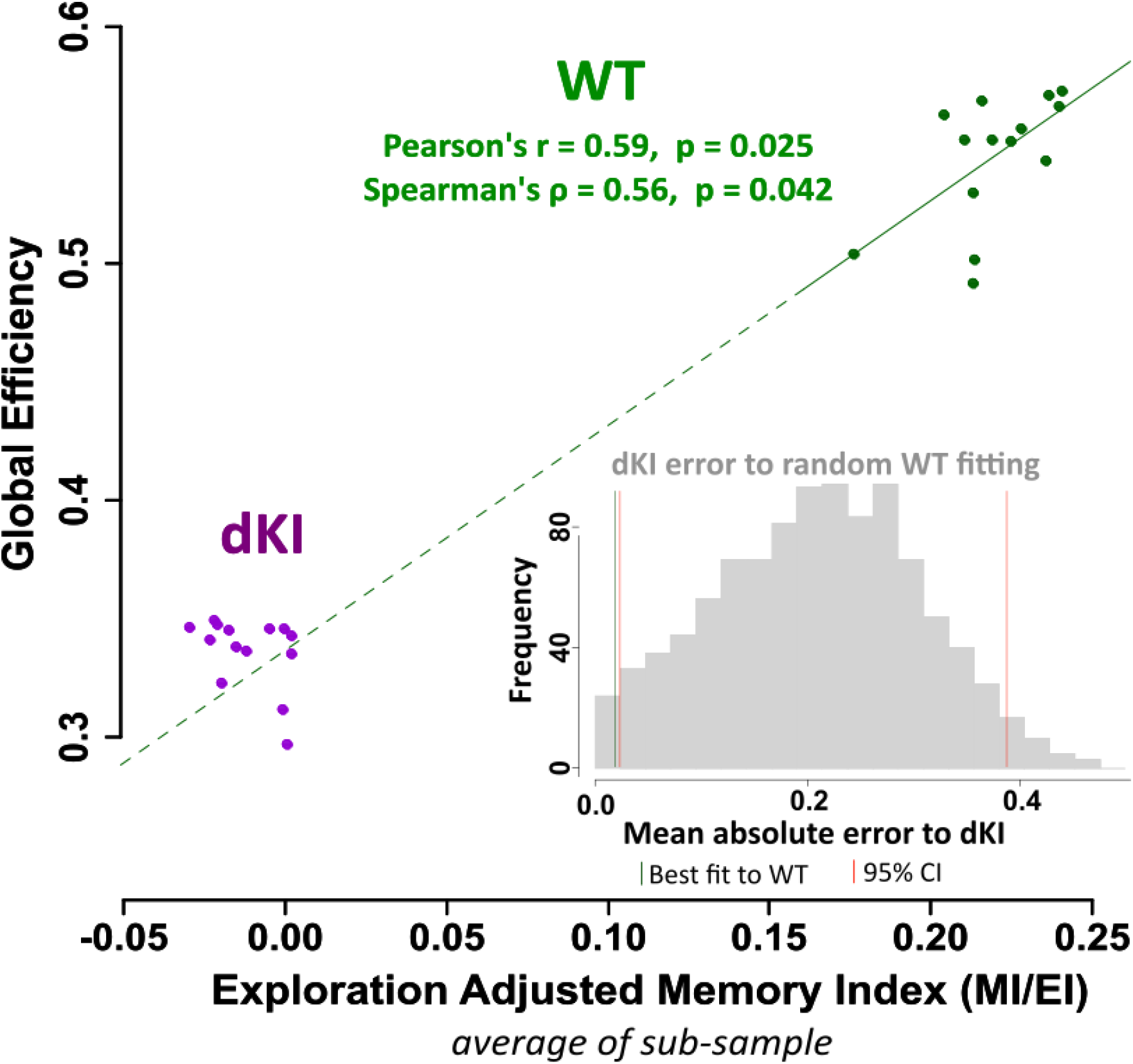
Association between network efficiency and exploration adjusted memory index ―MI/EI‖. Sub-samples were generated from each retention group by resampling n-1 mice without replacement. From each sub-sample, the average MI/EI and the network global efficiency were computed, where each sub-sample is depicted as point on the plot. The relationship between global efficiency and MI/EI was evaluated across only the WT sub-samples (solid green line), and a significant positive linear relationship was found. Interestingly, when this linear law across WT sub-samples was extrapolated down (dashed green line) to the reduced global efficiency of the dKI sub-samples, it could predict their reduction in MI/EI (the intersection of the dashed green line to the cloud of purple points). The prediction of dKI sub-samples MI/EI index based on extrapolating the law fitted on WT sub-samples yielded errors way smaller than what expected at chance level, as we could verify via comparison with a null distribution of prediction errors constructed from 1000 randomly fit models. See inset histogram: the green vertical line (dKI error to best fit of WT) lies to the left of the leftmost red vertical line (lower end of the null distribution 95% CI).

Moreover, we could show that extrapolating the best fit line for the WT ensemble to drops in global efficiency as large as the ones observed in the dKI ensemble also predicted the observed dKI decrease of the MI/EI behavioral performance index. This is illustrated in Figure 6 where the dashed continuation of the green line (the extrapolation of the best fit line to the WT ensemble) intersects the cloud of purple points (the ensemble of dKI sub-samples). The prediction of dKI sub-samples MI/EI index based on extrapolating the law fitted on WT sub-samples yielded errors way smaller than what expected at chance level, as we could verify via comparison with a null distribution of prediction errors constructed from 1000 randomly fit models to the WT ensemble. This is shown in the inset histogram of Figure 6, where the green vertical line (prediction error of best fit to WT) lies to the left of the leftmost red vertical line (lower end of the null distribution 95% CI).

This suggests that subjects with a high memory index but low total exploration time are thus more likely to have higher global efficiency. Contrary to memory index, there is no detectable significant difference in total object exploration time between the WT and dKI groups (Figure 6 - figure supplement 1C). Memory index is thus a more sensitive metric of subtle and complex behavioral changes, which are tracked by a decrease of global efficiency from WT to dKI. These results support reduced network integration as a potential factor for impairment of OP memory recall in the dKI.

## Discussion

The current necessity to learn more about initial steps of AD pathology prompted us to investigate brain network alterations associated to the encoding and retrieval phases of the first recognition memory paradigm affected in the late-onset *App*^*NL F*^/*MAPT* dKI mouse model of the disease (Saito et al, 2019). Here we show that the neuropathology expressed by 4-month-old dKI mice at the time of their earliest recognition deficit is reminiscent of an early preclinical stage. Quantification of c-Fos activation highlighted an abnormally increased activity in the entorhinal cortex and the claustrum, two entities known to be vulnerable to AD pathology. Finally, extensive analyses of network connectivity point to a disorganization of internal communication between and within the MTL and interconnected regions where the RSC seems to play a pivotal ―hub‖ role which could be disrupted in dKI mice.

### Object-place recognition deficit as an early marker of emerging AD neuropathology

In the battery of five recognition memory tasks, the object-place associative memory task was the first to detect a deficit in dKI mice at 4 months of age. This recognition deficit occurred 2 months before spatial pattern separation deficits and with intact long term object recognition memory up to 6 months of age. The object-place task seems to be highly sensitive to early amyloid pathology because it was also impaired at pre-plaque stages in more aggressive models of AD (Bonardi et al, 2016; Hamm et al., 2017). In TgCRND8 mice it was specifically associated with increased levels of β-CTF in the hippocampus (Bonardi et al, 2016; Hamm et al., 2017). Aβ levels of *App*^*NL F*^/*MAPT* dKI were not evaluated in this study, but the seminal work characterizing the parental single KI *App*^*NL F*^ mice already showed increased cortical levels of the neurotoxic species Aβ_42_ at the age of 2 months (Saito et al, 2014). This early accumulation of Aβ in 2-month *App*^*NL F*^ mice was associated with significant cell loss and hyperexcitation in the LEC, whereas the same alterations appeared several months later in the CA1 (Petrache et al., 2019). Here, we show for the first time evidence for early onset of AD pathology in 4-month-old dKI mice with specific increases in levels of β-CTF and phosphorylated tau labeled with AT270 antibodies in the MTC, but not in the dorsal hippocampus. These results suggest that early abnormal levels of tau phosphorylation are still restricted to the MTC at the age of 4 months. This is of particular importance because the PRC-EC pathway has long been seen as a primary site of neurodegenerative event and dysfunction that characterizes the earliest preclinical Braak stage I of AD tau deposition (Kahn et al, 2013: Braak and Braak, 1995; Hyman et al, 1984). Interestingly, significant worsening of neuropsychological markers of PRC and EC functionality were detected more than 10 years before AD diagnosis (Hirni et al, 2016). More recently, Yeung et al., 2018, showed that object-place recognition performance was predicted by anterolateral EC volume in the elderly. This result is in agreement with studies attributing a key role of the equivalent rodent LEC in the network supporting association of an object with a place or a scene (Deshmukh et al, 2012; Wilson et al, 2013a,b; Chao et al 2016). It must be noted that deficits in the pattern separation task in 6-month-old dKIs suggest subsequent development of functional alterations within the DG-CA3 region which is in agreement with studies in aMCI patients and transgenic mouse models of AD (Yassa et al, 2010; Zhu et al, 2017). All these findings strongly suggest that 4-month-old dKI mice recapitulate early impairment in recognition memory and specific vulnerability of the PRH-EC region to both amyloid and tau pathologies as found in preclinical stages of AD (Kahn et al, 2014).

### Distributed increase in c-Fos activation possibly reflecting spreading of pathological hyperactivity

We detected high level of c-Fos expression in the CLA and the LEC of dKI mice regardless of the test-phase suggesting that both structures would be hyperactivated during memory encoding and recall. On the other hand, the hyperactivation of the MEC restricted to the acquisition phase could be have been triggered by active encoding of the new environment. Well documented in mouse models of AD, early hyperactivity within the LEC appears as a consequence of local increase in amyloid pathology and it is considered as a major factor driving propagation of both Aβ and tau pathology to its main outputs especially the hippocampus (Xu et al., 2015; Nuriel et al., 2017; Rodriquez et al., 2020). The CLA has been less extensively studied, but a few studies show evidence of Aβ accumulation and neurodegeneration in this region (Qin et al, 2013; Ogomori et al., 1989; Gustafson et al., 1998). Interestingly, an early study in Alzheimer's patients found neuronal loss only within a CLA subregion strongly connected with the EC (Morys et al., 1996). Thus, EC may also propagate hyperactivity, exporting amyloid and tau pathology to the CLA as well as other limbic cortices in preclinical stages of AD (Bonthius et al., 2005). Note that an alternative hypothesis has been proposed with a central role of the CLA in the spreading of AD pathology in the brain (Avila and Perry, 2021). Being largely interconnected with most cortical areas (Wang et al, 2017), hyperactivity of CLA may initiate or aggravate spontaneous the cortical network hypersynchrony and task-induced hyperactivation found in elderly at risk and in early MCI (Mueller and Weiner, 2017; Corriveau-Lecavalier et al., 2020).

Early-stage hyperactivity has also been proposed to reflect compensatory mechanisms which would first help maintaining and then worsen cognitive function through the spreading of neuropathological processes and a deleterious impact on cognitive-related network organization and functioning (Corriveau-Lecavalier et al., 2020). There is some indication for a reorganization of network information flow in the dKI mice. We saw through the community organization of dKI functional networks during both phases, that the MEC and POC are more integrated with mPFC and DH communities (Figure 4C,D). This change in community allegiance may be a consequence of an initial hyperactive state within the EC, much like artificial stimulation of brain structures can enforce specific functional networks (Warren et al., 2019). In certain cases, this shift towards stronger outbound allegiance may help facilitate alternative communication between the MTC and interconnected regions, as seen with the maintenance of indirect information flow of the DG_sp during acquisition (Figure 5D,E). The apparition of compensatory hubs may be one way through which hyperactivity could initially help memory processing during the preclinical AD.

### mPFC and RSC are disrupted during object-place associative memory retention

The mPFC and RSC were the most heavily utilized regions in WT mice during retention phase, and direct and indirect information flow through these regions was significantly reduced in dKI mice (Figure 5F). The specificity of this dysfunction to the retention phase is not surprising as the mPFC and RSC are shown to be more involved in the retrieval and/or editing of memory traces rather than encoding *per se* (Mitchell et al., 2018). Lesioning the RSC disrupts object-place associative memory in rodents (Parron and Save., 2004), and communication between the mPFC-MTC is also shown to be essential (Chao et al., 2016; Hernandez et al., 2017). The RSC is proposed to play a pivotal ―hub‖ role in facilitating this communication as it exhibits strong structural connections with both the mPFC and the MTC (Sugar et al., 2011, Vann et al., 2009). In humans, the posterior cingulate cortex (PCC) and RSC, close equivalents to the rodent RSC (Lu et al., 2012, Stafford et al., 2014, Vogt and Paxinos, 2014), are shown to support structural connectivity between the mPFC and MTL, essential to associative memory networks and the DMN (Greicius et al., 2009, Miller et al., 2014). The hubness of the RSC during retrieval is clearly supported by the community organization of our functional networks, where it is well positioned to facilitate communication between the mPFC and MTL communities (Figure 4B). The severe loss in strength seen in the dysgranular RSC may therefore indicate an object-place recall-dependent roadblock for mPFC-MTL communication.

One question lies as to why the dysgranular RSC was more severely disrupted than the granular part. The dysgranular cortex is comparatively more connected to visual and sensory processing areas, making it ideal for processing and identifying local and distal cues for allocentric orientation. Lesioning the dysgranular RSC alone is enough to have rats shift from allocentric to egocentric strategies in a spatial memory task (Vann and Aggleton., 2005). The dysgranular RSC contains landmark dependent head direction cells, which may help orient the mice in the open field with respect to the objects (Jacob et al., 2016), and decreased strength of the dysgranular RSC may contribute to their dysfunction.

Our findings also corroborate those of human studies showing that the mPFC and the PCC/RSC are often disrupted in evaluations of functional and structural connectivity in aMCI (Catheline et al., 2010; Wang et al., 2013c). The especially severe disruption of the RSC reflects clinical observations of the human PCC as one of the most, if not the most, consistently disrupted regions in evaluations of resting state FC of aMCI (Badhwar et al., 2017; Eyler et al., 2019). The mPFC and the PCC/RSC are also the first regions to display amyloid deposition (Palmqvist et al., 2017). The PCC/RSC shows reduced connectivity in pre-plaque APOE ε4 carriers at risk for developing AD (Wang et al., 2013b; Jones et al., 2016), and very early increases in PCC amyloid deposition correlate with face-name associative memory deficits in subjective cognitive decline (Sanabria et al., 2017). Initial pathologies of amyloid in the mPFC/PCC/RSC and tau in the MTC may thus have a combined negative effect on associative memory through reducing memory-recall dependent FC.

### Decreased global efficiency during retention predicts memory deficits in dKI mice

In WT mice, we detected a positive relationship between global efficiency of the retention network and memory performance when taking into account a disruptive effect of total object exploration. The exploration disruption on memory driven FC can perhaps be explained by increased focus and a more conscious exploration of objects for mice with a low exploration index, in contrast to a more haphazard exploration of objects for mice with high exploration index. This may also indicate a disruptive effect of simultaneous sensory or motor network activation. Regardless, dKI mice presented no difference in object exploration as compared to the WT mice, making it an unlikely behavioral link to reduced global efficiency in these mice.

Global efficiency of c-Fos derived fear memory networks has already been shown as a reliable measure for predicting memory performance in mice (Vetere et al., 2017). Such findings are consistent with clinical observations linking increased global efficiency to better cognitive function (Li et al., 2009; Stanley et al., 2015). The relationship across the WT group can be extrapolated down to predict memory performance in the dKI. This provides further evidence that their drop in memory performance is directly linked to deficient information transfer of the memory retention network. There is increasing evidence that memory loss in AD pathology is related to dysfunctional recall, as it can be rescued by aiding retrieval processes (i.e., activating the ―silent‖ engram; Roy et al., 2016, Perusini et al., 2017). Perhaps early associative memory deficits of dKI mice could be rescued by activating ―silent‖ inefficient functional networks. In humans, noninvasive brain stimulation has been extensively studied as a potential therapeutic tool for AD on regions such as the dorsolateral mPFC with mixed results (Weiler et al., 2020). However, electromagnetic stimulation of the parietal cortex was shown to support associative memory by increasing memory-retrieval dependent FC strength of an associated parietal-RSC-hippocampal network (Wang et al., 2014; Warren et al., 2019). Taken together with our study, these results suggest that the PCC/RSC, or more laterally accessible and functionally similar regions such as the parietal cortex, may be more effective targets for improving associative memory in AD.

### Contrast with resting state fMRI in other mouse models of AD

Contrary to our results, studies that evaluate resting state networks in pre-plaque or early-NFT mouse models of AD predominantly find cortical and hippocampal-cortical hyperconnectivity, while hypoconnectivity appears at later stages (Asaad and Lee, 2018). This suggests that perturbations in resting state FC may predict memory FC perturbations, but do not directly mirror them. Moreover, the mouse models used to evaluate pre-aggregate resting state FC presented either amyloid pathology (Shah et al., 2016; Shah et al., 2018) or tau pathology (Degiorgis et al., 2020) but not both concurrently. It has been shown with combined resting state fMRI and PET imaging in humans that increased Aβ alone is associated with hyperconnectivity of the DMN while combined Aβ and Tau pathologies reveal hypoconnectivity (Schultz et al., 2017). Whether these contrasting results reflect differences between―resting state vs memory driven FC‖ or differences in ―amyloid/tau pathological staging‖ will be more thoroughly understood once resting state fMRI is directly measured in young dKI mice.

In conclusion, our results suggest that the present dKI model was caught at the very beginning of its neuropathology as it was restricted to the MTC region, leaving the dorsal hippocampus quite preserved. The local MTC pathology of these mice was associated with EC and CLA hyperactivity which would most likely spread towards densely interconnected regions such as the mPFC, the DH and the RSC. Retrieval dependent communication between cingulate areas and the MTL was disrupted, and can be potentially linked to reduced dysgranular RSC hub strength. The similarity between our findings in the dKI model and those reported in the earliest stages of the disease suggests that the *App*^*NL-F*^ version of the dKI model has a high potential for generating new discoveries on the earliest stage of AD.

## Methods

### Animals

The *App*^*NL-F*^/*MAPT* double knock-in (dKI) mice were produced through crossing homozygotes of two single knock-in (KI) mouse lines for the humanized *App*^*NL-F*^ and *MAPT* genes, and then through the crossing of the resulting doubly heterozygote mice to obtain dKI and non-knock-in WT line founders. After three generation of homozygous breeding, the dKI mouse line was backcrossed with C57BL/6J mice (Janvier Laboratories, Le Genest Saint Isle, France) in order to limit genetic drift between dKI mice and their WT controls. The *App*^*NL-F*^ gene contains a humanized Aβ fragment with Beyreuther/Iberian and Swedish FAD mutations, and the human *MAPT* gene expresses all 6 isoforms of tau found in humans. Both single KI mouse lines were produced by T Saido and T Saito (RIKEN Brain Science Institute, JAPAN) and sent to us by the RIKEN BioResource Center. Mice were group-housed with food and water ad libitum, nesting material, and additional food pellets on bedding to promote natural behavioral patterns. The animal room was under controlled temperature (23 °C ± 1°C) and a 12/12-hour light/dark cycle (lights on at 8.00 AM). Procedures were in compliance with rules of the European Community Council Directive 2010-63 and French Department of Agriculture Directive 2013-118 and approved by the local review board (CREMEAS: APAFIS#9848). Animal facilities were approved for animal experimentation (H 67-482-13).

### Behavioral Testing

Behavioral testing took place during the light phase. Mice were single-housed for 1 week before testing. Spontaneous object exploration tests were carried out in an open field (100cmx100cmx50cm, Ugo Basile, Italy) with dark grey acrylic walls and a grey metal floor. The open field was evenly illuminated by three indirect halogen lights (open field center, 15 lux), and a radio gave background noise from 1.5m away (open field center, 45 ± 5 dB). Nine different sets of objects were used: two for the habituation phases, two for the long-term novel object recognition task, two for the OP task, two for the short-term novel object recognition task, and one for the object location task. These objects differed in size (10 to 20 cm), material (metal, glass, or plastic), shape, and color. Each object was available in duplicate or triplicate. Ethanol (30%) was used to clean the objects and the open field between each trial. Object exploration time was recorded and defined as the nose pointing toward the object within 2 cm. Gnawing and climbing of objects were not counted as exploration time.

### Habituation

Before testing, all mice received two days of habituation. On the first day, they were given a habituation trial of 10 min with two identical objects placed in the open field. On the second day they were given two 10 min trials with a different set of two identical objects, with the trials separated by an inter trial interval (ITI) of 5‘ that the mice spent in their home cage. For the OP task with a 3-hour ITI, the second day habituation procedure was applied with an ITI of 3 hours.

### Preliminary testing cohorts

Mice were tested at 2 months (WT n = 10, dKI n = 10), 4 months (WT n = 12, dKI n = 9) and at 6 months (WT n = 8, dKI n = 11). An additional cohort of (WT n = 3, dKI n = 1) was tested at 2 months in object-place association as the initial WT group did not reach significance above 0. For pattern separation one two month old WT mouse was removed due to the wrong objects accidentally being placed during the retention phase.

### Short term, pattern separation and long-term novel object recognition

Mice were tested at 4 months (WT n = 11, dKI n = 11) in the short term OR task. One WT mouse was removed from the analysis due to being dropped before the task. Mice explored two identical objects during a 10-min acquisition trial, and were returned to their home cage for an ITI of 5 min (short term, pattern separation) or 24 hours (long term). Thereafter, mice were given a 10-min retention trial, where one of the familiar objects was replaced by an unfamiliar new one. For the pattern separation task, objects were made of legos, the novel object had the same composition of colored lego blocks but a different pattern. Exploration of the objects was recorded during the 6 minutes (4 minutes for pattern separation) following the initial exploration of object for each mouse. The memory index was calculated as:

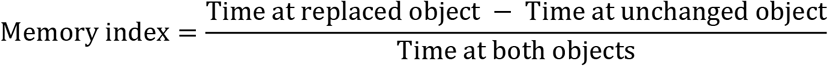

### Short term object-place association

To validate the potential deficit seen during preliminary phenotyping, an additional cohort (WT n = 11, dKI n = 11) was tested in the object-place association task. Mice explored two different objects during a 10-min acquisition trial, and were returned to their home cage for an ITI of 5 minutes. During the 10-min retention trial, one of the objects was replaced by a copy of the other. The mice had to detect the mismatch between one object and its actual location in the open field. Exploration of the objects was recorded during the 4 minutes following initiation of object exploration episode for each mouse. The memory index was calculated as with the long-term object recognition task.

### Short term novel object location

Mice were tested at 4 months (WT n = 11, dKI n =11). Mice explored two identical objects during a 10-min acquisition trial, and were returned to their home cage for an ITI of 5 minutes. During a 10-min retention trial, one of the objects was moved 55 cm from its original position for the object location task. Exploration of the objects was evaluated during the 6 minutes following the initiation of object exploration for each mouse. During both phases, all objects were placed equidistant from the walls. The memory index was calculated as:

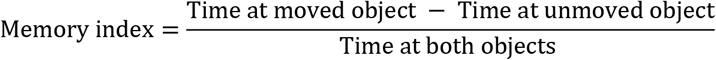

### Western Blotting

Four-month-old male mice were sacrificed by cervical dislocation (n=8 per group), and their brains were carefully dissected on ice. Hippocampi and medial temporal cortex were quickly removed, frozen in liquid nitrogen and then stored at −80°C until their use. Tissues were homogenized in 10 volumes of ice-cold radioimmunoprecipitation assay buffer containing protease inhibitor cocktail (Sigma-Aldrich, St. Louis, Missouri), phosphatase inhibitor cocktail (PhosStop, Roche Life Science, Penzberg, Germany), and 1 mM phenylmethylsulfonyl fluoride (Sigma-Aldrich). After centrifugation at 20,000g for 20 min at 4°C, supernatants were aliquoted for immunoblot analysis. Brains extracts of Tg2576 and Thy-tau22 mice were homogeneized and separated in parallel to our samples as a positive control for APP proteins and cleaved fragments and for tau proteins, respectively. Protein concentration was measured using the Bio-Rad Protein Assay (Bio-Rad, Hercules, California). Thirty or 20 microgramms were respectively loaded on 4-20% precast gel (Mini-Protean TGX precast gels, Bio-Rad) for APP and tau proteins. After electrophoresis and transfer to nitrocellulose membranes using the Trans-Blot Turbo System (Bio-Rad), membranes were incubated with 5% skimmed milk for 1 hour at room temperature and then with primary antibodies diluted in 2% bovine serum albumin (Sigma-Aldrich) in tris-buffered saline 0.05% Tween 20 (Sigma-Aldrich) overnight at 4°C. After washes, membranes were incubated with anti-mouse or anti-rabbit immunoglobulins conjugated to horseradish peroxidase (Jackson Immunoresearch, West Grove, Pennsylvania) for development with enhanced ECL chemiluminescence detection kit (Thermo Fisher Scientific, Waltham, Massachusetts). After detection, all membranes were re-probed with anti-actin antibody for normalization of total protein. The primary antibodies used were the rabbit polyclonal anti-APP, C terminus (Sigma-Aldrich), the rabbit polyclonal anti-tau (B19, generously gifted by JP Brion, ULB, Belgium), the mouse monoclonal anti-phopho-Tau Thr181 (AT270, ThermoScientific), the mouse monoclonal anti-actin (Sigma-Aldrich) and the rabbit polyclonal anti-actin (Sigma-Aldrich). The secondary antibodies used were the peroxidase-conjugated AffiniPure goat anti-mouse and goat anti-rabbit (Jackson Immunoresearch). The quantification of the band intensity acquired with the ChemiDoc Imaging system (Bio-Rad) was performed by densitometry analysis using the ImageJ program. For each mouse, the phosphorylation degree was calculated as the ratio of total phosphorylated Thr181 tau proteins on total tau proteins. The ratio of APP-cleaved fragments was calculated as the total of APP β-CTF on α-CTF fragments.

### Object-place association with a 3-hour ITI for ex-vivo imaging

The same protocol as with the short-term object-place task was followed, except with an ITI of 3-hours. The ―Acquisition‖ set of WT (N=12) and dKI (N=12) mice were only tested in the acquisition phase. The―Retention‖ set of WT (N=14) and dKI (N=14) mice were tested for the entire task, acquisition and retention phases included. At the end of testing mice were left in their home cage in a quiet dim room (9 lux, 35 ± 5 dB) next to the testing room for 90 minutes, after which they were taken for brain perfusion and removal.

### Perfusion and tissue preparation

Mice were killed with an overdose of sodium pentobarbital (105 mg/kg intraperitoneally) and transcardially perfused in 0.1% heparin phosphate-buffered saline (PBS) and 4% paraformaldehyde [PFA; in phosphate buffer (PB) pH7.4; 4°C]. Brains were removed, postfixed in a 4% PFA solution for 24h, and cryoprotected in a saccharose solution (20% in PB, 0.1 M; pH 7.4; 4°C) for 48 hours before being frozen with isopentane (−40°C) and subsequently stored at −80°C. Forty μm coronal cryostat sections were cut from the anterior to the posterior of the brain. For the medial prefrontal cortex (mPFC), claustrum (CLA), dorsal hippocampus (DH), medial entorhinal cortex (MEC), and postrhinal cortex (POC) every 4^th^ section created a set (spacing of 160 μm), and for the retrosplenial cortex (RSC), lateral entorhinal cortex (LEC), and perirhinal cortex (PRC) every 6^th^ section (spacing of 240 μm).

### Immunohistochemistry

Brain sections were given three 10 min washes in 0.1 M phosphate-buffered saline (PBS), followed by a 30 min incubation in 1% H_2_O_2_. They were washed for 5 min with ultra-pure water and again three times for 10 min in 0.1 M PBS. This was followed by a 45 min blocking incubation in a 5% natal goat serum (NGS) diluted in a ―diluent‖ solution consisting of 0.1 M PBS, and 0.5% triton. The sections were incubated at room temperature for 2 days in a 1/15000 dilution of rabbit-anti-cFos (Synaptic Systems, Göttingen, Germany) primary antibody in diluent containing 2% NGS. After 2 days the sections were first given two 10 min washes in 0.1 M PBS, and then incubated for 2 hours at room temperature in a 1/500 dilution of biotinylated mouse-anti-rabbit (Vector Laboratories, Burlingame, California) secondary antibody in diluent containing 2% NGS. The sections were given two 10 min washes in PBS, followed by a 45 min incubation in the avidin/biotin (Vector Laboratories) solution. This was followed by three 10 min washes in 0.1 M PBS and a 10 min wash in phosphate buffer. The sections were finally revealed with a 10 min incubation in 3,3-diaminobenzidine (Vector Laboratories). Images of whole brain sections were taken at 20x magnification using a Hamamatsu NanoZoomer S60 digital slide scanner (Hamamatsu Photonics K.K., Hamamatsu City, Japan) for offline quantification of c-Fos expression.

### c-Fos Imaging

Neuronal activation is associated with increases in intracellular calcium levels, which in turn leads to the rapid up-regulation of immediate early genes such as c-fos. The quantification of c-Fos protein levels can thus be used to derive a measure of neuronal activity (Tischmeyer and Grimm, 1999). c-Fos expression was analyzed in 22 regions of interest (ROIs) (see Table S1), including sub-regions of the PFC, CLA, DH, RSC, PRC, POC and EC. Image processing was done using ImageJ (National Institute of Health, Bethesda, MD). ROIs were anatomically defined according to the atlas of Franklin and Paxinos (2008). For c-Fos quantification, the images were transformed into 8-bit grayscale. A grayscale threshold was set at a consistent level for each region by an experimenter blind to group condition. Only c-Fos positive nuclei with a grayscale intensity below the threshold and an area between 25–300 μm^2^ were counted. At least three brain sections were processed per ROI. Mean c-Fos density was calculated for each ROI as the quantity of c-Fos marked nuclei per mm^2^, normalized to the WT-Acquisition group. ROIs were grouped into region subfields whenever anatomically and functionally justified. These groupings reduce the number of comparisons and, thereby, restrict Type 1 errors. Three-way ANOVAs compared test-phases (acquisition or retention), genotypes (WT or dKI) and subregions for each subfield. When an interaction was significant, the simple effects were examined.

### Functional Connectivity

From the c-Fos signals, functional connectivity (FC) was assessed by computing the between subject inter-regional spearman correlations for each Genotype-Phase group. Spearman correlations, rather than Pearson correlations, were used to account for potential outlier effects and the relatively small sample sizes. Correlation matrices were used to visualize all possible pairwise inter-regional correlations within each group. To assess global FC strength the mean r was calculated with retained, near-zeroed, and absolute valued negative correlations respectively. In the near-zeroed case, negative correlations were reduced to a value of .006, the smallest positive correlation observed.

### Generating functional networks as fully connected weighted graphs

From each correlation matrix, a functional network was generated as a fully connected weighted graph. The edges weights of the graph reflect inter-regional spearman correlation strengths and the nodes reflect regions. Negative correlations can be interesting, but complicate considerably graph analyses as various algorithm variants exist to handle them (e.g. in community detection) and there is no obvious criterion to choose one variant over others. For the sake of clarity, we thus treated negative correlations in most analyses (unless explicitly mentioned) as near-zero positive value of correlation (minimum edge weight of 0.006, see above). In efficiency analyses, this corresponds to interpreting negative correlations as open but difficult paths for information transfer. All graph construction and graph analysis were done through the igraph (Csardi G and Nepusz T, 2006) package on R (R Core Team, 2017).

### Bootstrapping confidence intervals

For all network metrics, confidence intervals were computed through bootstrapping. This involves resampling subjects with replacement 1000 times, each time regenerating a functional network, then recalculating the estimate of interest. The 95% quartile of the bootstrap distribution was taken as the 95% confidence interval. Confidence intervals for the difference were used to test for differences between genotype groups (Wright et al., 2011). Groups were considered different to a P<.05 if the 95 % confidence interval for the difference ≥ 0, and to a P<.01 if the 99 % confidence interval for the difference ≥ 0.

### Community Analysis

Networks with high modularity, ***Q***, have strong connections between the nodes within communities and relatively weaker connections between nodes of different communities.

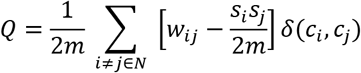

where *N* denotes the set of all nodes in the network, *w*(*i*, *j*) denotes the total number of edges in the network, denotes the edge weight between a node *i* and another node *j*, *s* denotes the sum of a node‘s edge weights, *c* denotes the community to which a node belongs, and *δ*(*c*_*i*_, *c*_*j*_) indicates if the compared nodes are in the same community ( *δ*(*c*_*i*_, *c*_*j*_) is 1 if *c*_*i*_ = *c*_*j*_, and 0 otherwise). Communities were detected in each bootstrap through finding the maximum modularity across all possible community partitions. This modularity maximization computation was done through the *cluster_optimal* function of the igraph package. This transforms modularity maximization into an integer programming problem, and calls the GNU Linear Programming Kit (GLPK) to solve that. See Brandes et al, 2008 for more details. This computationally expensive detection method of evaluating all possible community partitions for modularity maximization (in contrast to less expensive methods that infer modularity, as through greedy optimization, i.e. the Louvain method), could be permitted due to the relatively small number of nodes in our network. Allegiance matrices were used to assess community stability across bootstraps, by depicting the percentage of bootstraps (n = 1000) that contain any given pair of regions within the same community.

### Information flow

Nodal strength, ***s***, is traditionally calculated as the sum of a node‘s edge weights. In a fully connected network this is directly proportional to the average of a node‘s edge weights. The average, ***s***_***average***_, was used in our case for better comparison with nodal efficiency.

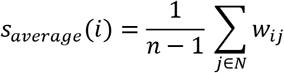

where *N* denotes the set of all nodes in the network, *n* denotes the total number of nodes in the network, and *w*_*ij*_ denotes the edge weight between a node *i* and another node *j*.

For efficiency metrics, edge lengths were first computed as inverted edge weights. Nodal efficiency, ***E***_*nodal*_, was calculated as the average inverse shortest path length between a region and all other regions of the network.

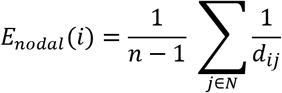

where *d*_*ij*_ denotes the length of the shortest path (lowest sum of edge lengths) between a node *i* and another node *j*. Global efficiency, ***E***_*global*_, was calculated as the average inverse shortest path length of the network.

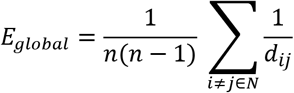

### Assessing the impact of global efficiency on memory deficits

Subsamples were generated from each retention group by resampling n-1 mice without replacement. This procedure generates a set of subsamples of size n-1, where the number of subsamples is the number of mice, n. From each subsample, the average memory index and the resulting network global efficiency were computed. Correlation significance between memory index and global efficiency was computed through both Pearson and Spearman correlation coefficients. To compute the exploration adjusted memory index, first the exploration index was defined for each mouse as total time exploring both objects normalized to the group average:

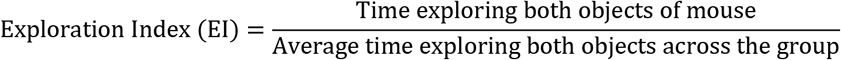

The exploration adjusted memory index was then defined as the memory index divided by the exploration index.

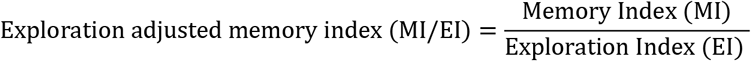

To assess the significance of the intersection of the WT linear fit with the dKI subsample population, 1000 randomly fit linear models to the WT were generated by permuting the y labels of the WT subsample population and recalculating the best fit line each time. The mean absolute error to the dKI subsamples was calculated for each random model and their distribution is presented as a histogram. The 95% confidence interval was considered as the 95% quartile of the distribution, and compared to the dKI mean absolute error to the original WT fit.

---

**Figure 1 - figure supplement 1.**
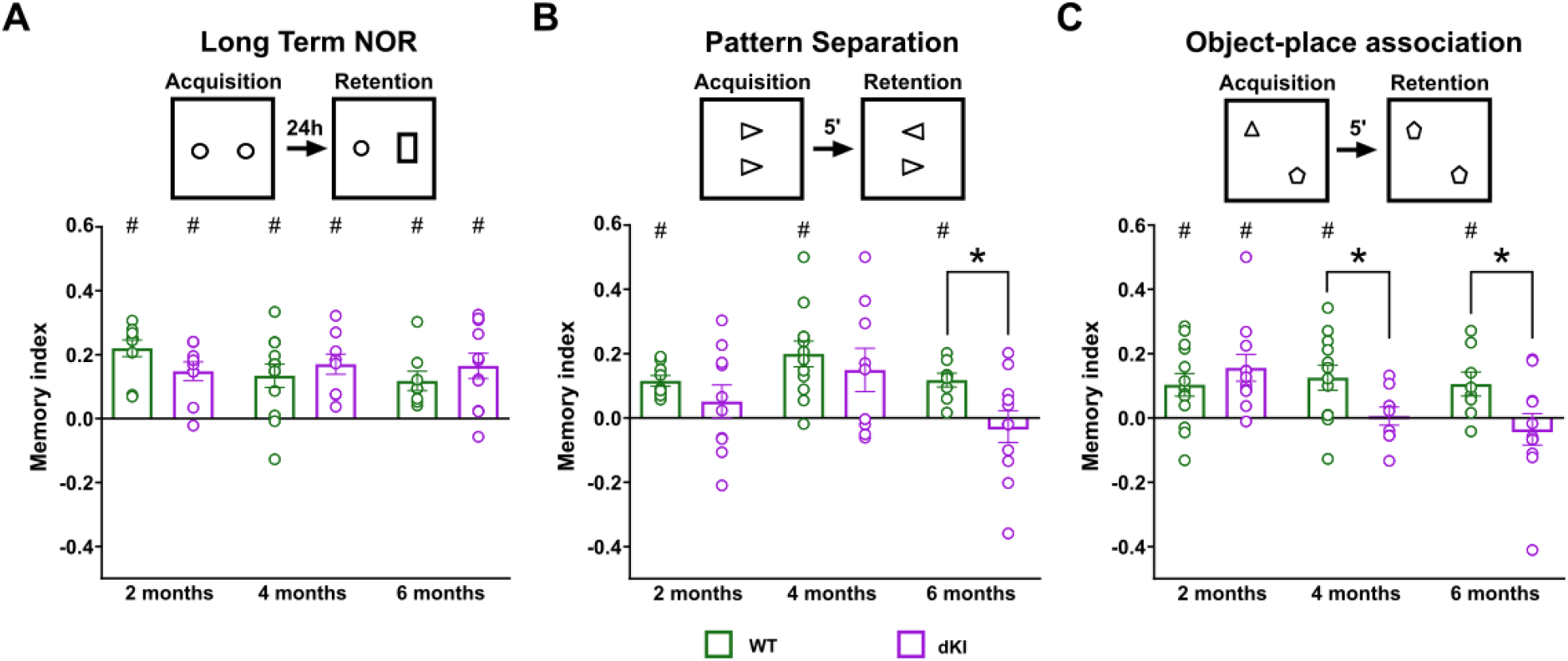
Preliminary phenotyping of young dKI mice with sensitive recognition memory paradigms. Mice were tested at 2 months (WT n = 10, dKI n = 10), 4 months (WT n = 12, dKI n = 9) and at 6 months (WT n = 8, dKI n = 11). An additional cohort of (WT n = 3, dKI n = 1) was tested at 2 months in object-place association as the initial group did not reach significance above 0. (A) In the long term object recognition task, dKI mice were completely unaffected. (B) In the pattern separation task the dKI mice were highly variable making it so that they were not significant against chance at any age (one sample t-test, against chance: all dKI ts<.996, ps>0.055). However, between group significance was only obtained at 6 months of age (WT vs. dKI: t(17) = 2.364, P = 0.030). This variability may represent a variety of factors including perhaps affected DG neurogenesis (Scopa et al., 2020). (C) The earliest significant deficit in dKI mice appeared at 4 months in the object-place association task, a deficit which was conserved at 6 months (two sample t-test, WT_4mo vs. dKI_4mo: t(20) = 2.325, P = 0.0313; one sample t-test, against chance: dKI t(8) = 0.2086, P = 0.8399; two sample t-test, WT_6mo vs. dKI_6mo: t(17) = 2.132, P = 0.0479; one sample t-test, against chance: dKI t(10) = 0.7196, P = 0.4883). Although the variability seen during pattern separation may be interesting, we chose to continue our study with the more robust object-place deficit. Bar graphs represent the mean density (± SEM). difference between genotypes *p<.05 (two sample t-test); difference from chance, #p<.05 (one sample t-test). No multiple comparisons corrections were performed for these preliminary comparisons.

**Figure 1 - figure supplement 2.**
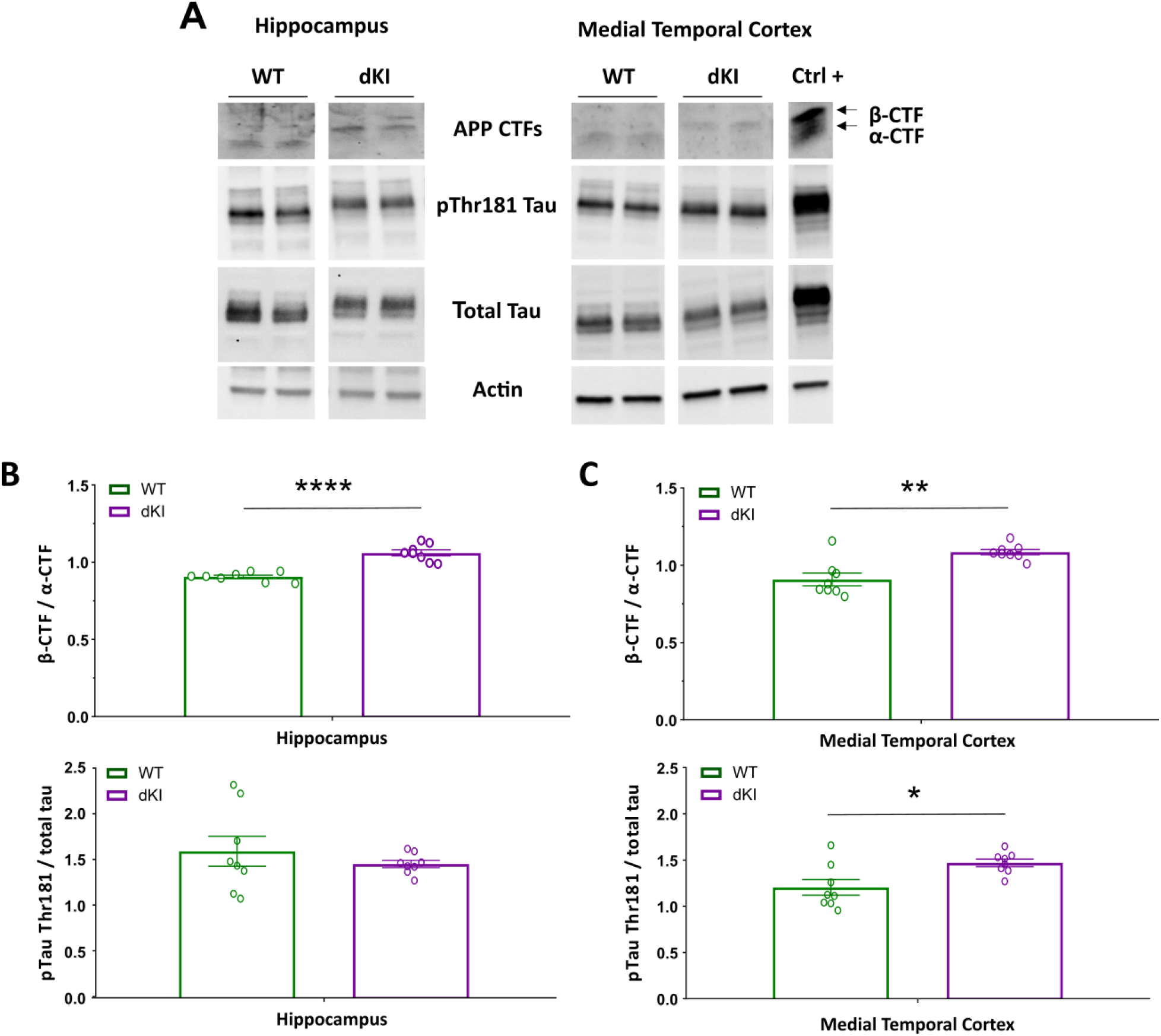
Early stage of Alzheimer-like disease in 4 month old male dKI mice. (A) Representative blots of APP-cleaved fragments, phosphorylated tau proteins on the threonine 181 site and of total tau proteins in the hippocampus and medial temporal cortex from WT and dKI mice. Analysis of the ratio of the β-CTF on α-CTF fragments and the degree of tau phosphorylation on Thr181 in the (B) hippocampus and in the (C) medial temporal cortex. Results showed that dKI mice expressed mainly the β-CTF whereas the WT mice expressed mainly the α-CTF and the β-CTF/α-CTF ratio increased in both brain regions. Only the MTC displayed an increase of phosphorylated tau proteins. Bar graphs represent the mean density (± SEM). * p<0.05, ** p<0,01, *** p<0,001 and **** p<0,0001 (two sample t-test).

**Figure 6-figure supplement 1.**
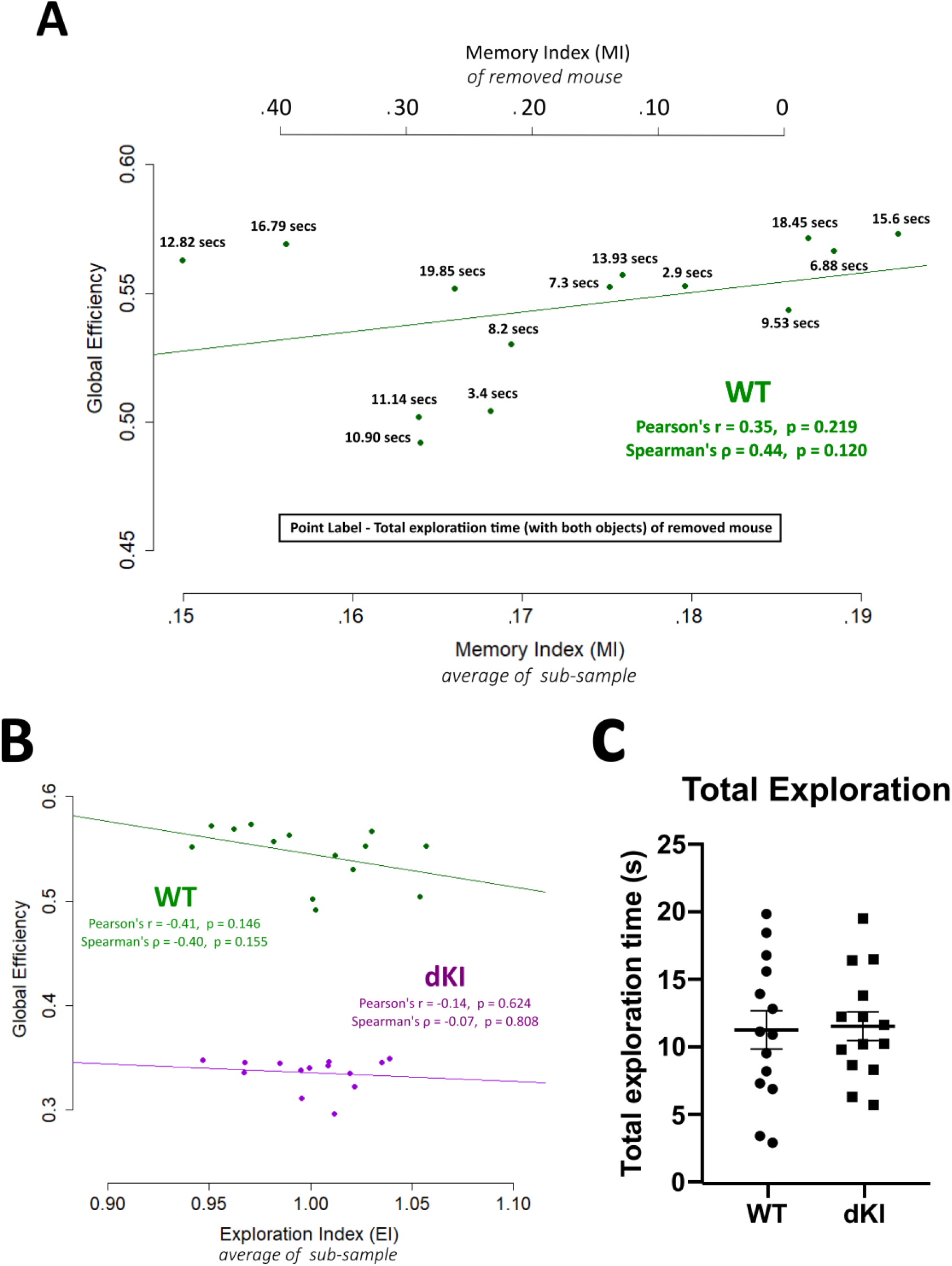
Associations of memory index ―MI‖ and exploration index ―EI‖ to network efficiency. (A) There was no relationship found between global efficiency and memory index. However, we noticed that of the mice with high memory index, those with greater exploration times (total time exploring both objects) appeared to have a weaker contribution to global efficiency. To take into account an exploration disruption effect on memory related FC, an exploration adjusted memory index was computed as the memory index divided by the exploration index (MI/EI), where the exploration index (EI) is the exploration time of the mouse normalized to the group. (B) We verified that there was no direct relationship found between global efficiency and EI. (C) There was also no difference in total exploration between WT and dKI mice.

## Abbreviations

AD: Alzheimer‘s disease
CLA: claustrum
DH: dorsal hippocampus
dKI: *App*^*NL-F*^/*MAPT* double knock-in mouse
FC: functional connectivity
mPFC: medial prefontal cortex
MTC: medial temporal cortex
MTL: medial temporal lobe
OP: object-place association
PS: Pattern Separation
RSC: retrosplenial cortex

## Acknowledgements

Université de Strasbourg, Centre National, de la Recherche Scientifique, complementary thesis support from the FRM-ALZ201912009643, Anne Pereira de Vasconcelos for advices on experimental design, Laura Durieux for analysis discussion, Dominique Massotte for the nanozoomer, Aminé Isik for mouse genotyping and breeding, Olivier Bildstein for care to our mice, JP Brion for the B19 anti-body.

## References

Asaad M, Lee JH. 2018. A guide to using functional magnetic resonance imaging to study Alzheimer‘s disease in animal models. Dis Model Mech 11. doi:10.1242/dmm.031724

Avila J, Perry G. 2021. A Multilevel View of the Development of Alzheimer‘s Disease. Neuroscience 457:283–293. doi:10.1016/j.neuroscience.2020.11.015

Bai L, Zhang M, Chen S, Ai L, Xu M, Wang D, Wang Fei, Liu L, Wang Fang, Lao L. 2013. Characterizing acupuncture de qi in mild cognitive impairment: relations with small-world efficiency of functional brain networks. Evid Based Complement Alternat Med 2013:304804. doi:10.1155/2013/304804

Bassett DS, Bullmore E. 2006. Small-world brain networks. Neuroscientist 12:512–523. doi:10.1177/1073858406293182

Bassett DS, Yang M, Wymbs NF, Grafton ST. 2015. Learning-induced autonomy of sensorimotor systems. Nat Neurosci 18:744–751. doi:10.1038/nn.3993

Bergmann E, Zur G, Bershadsky G, Kahn I. 2016. The Organization of Mouse and Human Cortico-Hippocampal Networks Estimated by Intrinsic Functional Connectivity. Cereb Cortex 26:4497–4512. doi:10.1093/cercor/bhw327

Bonardi C, Pardon M-C, Armstrong P. 2016. Deficits in object-in-place but not relative recency performance in the APPswe/PS1dE9 mouse model of Alzheimer‘s disease: Implications for object recognition. Behavioural Brain Research 313:71–81. doi:10.1016/j.bbr.2016.07.008

Braak H, Braak E. 1995. Staging of Alzheimer‘s disease-related neurofibrillary changes. Neurobiol Aging 16:271–278; discussion 278-284. doi:10.1016/0197-4580(95)00021-6

Brandes U, Delling D, Gaertler M, Gorke R, Hoefer M, Nikoloski Z, Wagner D. 2008. On Modularity Clustering. IEEE Trans Knowl Data Eng 20:172–188. doi:10.1109/TKDE.2007.190689

Catheline G, Periot O, Amirault M, Braun M, Dartigues J-F, Auriacombe S, Allard M. 2010. Distinctive alterations of the cingulum bundle during aging and Alzheimer‘s disease. Neurobiology of Aging 31:1582–1592. doi:10.1016/j.neurobiolaging.2008.08.012

Caviezel MP, Reichert CF, Sadeghi Bahmani D, Linnemann C, Liechti C, Bieri O, Borgwardt S, Leyhe T, Melcher T. 2020. The Neural Mechanisms of Associative Memory Revisited: fMRI Evidence from Implicit Contingency Learning. Front Psychiatry 10. doi:10.3389/fpsyt.2019.01002

Chao OY, Huston JP, Li J-S, Wang A-L, Silva MA de S. 2016. The medial prefrontal cortex—lateral entorhinal cortex circuit is essential for episodic-like memory and associative object-recognition. Hippocampus 26:633–645. doi:10.1002/hipo.22547

Chen G, Xu T, Yan Y, Zhou Y, Jiang Y, Melcher K, Xu HE. 2017. Amyloid beta: structure, biology and structure-based therapeutic development. Acta Pharmacologica Sinica 38:1205–1235. doi:10.1038/aps.2017.28

Corriveau-Lecavalier N, Duchesne S, Gauthier S, Hudon C, Kergoat M, Mellah S, Belleville S, for the Consortium for the Early Identification of Alzheimer‘s Disease-Quebec (CIMA-Q). 2020. A quadratic function of activation in individuals at risk of Alzheimer‘s disease. Alzheimer’s & Dementia: Diagnosis, Assessment & Disease Monitoring 12. doi:10.1002/dad2.12139

Csardi G, Nepusz T. 2006. The igraph software package for complex network research. InterJournal Complex Systems:1695. doi:https://igraph.org

Degiorgis L, Karatas M, Sourty M, Faivre E, Lamy J, Noblet V, Bienert T, Reisert M, von Elverfeldt D, Buée L, Blum D, Boutillier A-L, Armspach J-P, Blanc F, Harsan L-A. 2020. Brain network remodelling reflects tau-related pathology prior to memory deficits in Thy-Tau22 mice. Brain. doi:10.1093/brain/awaa312

Deshmukh SS, Johnson JL, Knierim JJ. 2012. Perirhinal cortex represents nonspatial, but not spatial, information in rats foraging in the presence of objects: Comparison with lateral entorhinal cortex. Hippocampus 22:2045–2058. doi:10.1002/hipo.22046

DiCiccio T. J. and B. Efron. (1996). Bootstrap Confidence Intervals. Statistical Science. 11(3): 189–212. https://doi.org/10.1214/ss/1032280214

Drzezga A, Becker JA, Van Dijk KRA, Sreenivasan A, Talukdar T, Sullivan C, Schultz AP, Sepulcre J, Putcha D, Greve D, Johnson KA, Sperling RA. 2011. Neuronal dysfunction and disconnection of cortical hubs in non-demented subjects with elevated amyloid burden. Brain 134:1635–1646. doi:10.1093/brain/awr066

Etter G, van der Veldt S, Manseau F, Zarrinkoub I, Trillaud-Doppia E, Williams S. 2019. Optogenetic gamma stimulation rescues memory impairments in an Alzheimer‘s disease mouse model. Nature Communications 10:5322. doi:10.1038/s41467-019-13260-9

Eyler LT, Elman JA, Hatton SN, Gough S, Mischel AK, Hagler DJ, Franz CE, Docherty A, Fennema-Notestine C, Gillespie N, Gustavson D, Lyons MJ, Neale MC, Panizzon MS, Dale AM, Kremen WS. 2019. Resting State Abnormalities of the Default Mode Network in Mild Cognitive Impairment: A Systematic Review and Meta-Analysis. J Alzheimers Dis 70:107–120. doi:10.3233/JAD-180847

Franklin K, Paxinos G. 2008. The mouse brain in stereotaxic coordinates. San Diego: Academic Press.

Filippi M, van den Heuvel MP, Fornito A, He Y, Hulshoff Pol HE, Agosta F, Comi G, Rocca MA. 2013. Assessment of system dysfunction in the brain through MRI-based connectomics. The Lancet Neurology 12:1189–1199. doi:10.1016/S1474-4422(13)70144-3

Gardini S, Venneri A, Sambataro F, Cuetos F, Fasano F, Marchi M, Crisi G, Caffarra P. 2015. Increased functional connectivity in the default mode network in mild cognitive impairment: a maladaptive compensatory mechanism associated with poor semantic memory performance. J Alzheimers Dis 45:457–470. doi:10.3233/JAD-142547

Grajski KA, Bressler SL. 2019. Differential medial temporal lobe and default-mode network functional connectivity and morphometric changes in Alzheimer‘s disease. NeuroImage: Clinical 23:101860. doi:10.1016/j.nicl.2019.101860

Granholm E, Butters N. 1988. Associative encoding and retrieval in Alzheimer‘s and Huntington‘s disease. Brain Cogn 7:335–347. doi:10.1016/0278-2626(88)90007-3

Greicius MD, Supekar K, Menon V, Dougherty RF. 2009. Resting-State Functional Connectivity Reflects Structural Connectivity in the Default Mode Network. Cereb Cortex 19:72–78. doi:10.1093/cercor/bhn059

Guo Q, Li H, Cole AL, Hur J-Y, Li Y, Zheng H. 2013. Modeling Alzheimer‘s Disease in Mouse without Mutant Protein Overexpression: Cooperative and Independent Effects of Aβ and Tau. PLOS ONE 8:e80706. doi:10.1371/journal.pone.0080706

Gustafson L, Brun A, Englund E, Hagnell O, Nilsson K, Stensmyr M, Öhlin A-K, Abrahamson M. 1998. A 50-year perspective of a family with chromosome-14-linked Alzheimer‘s disease. Hum Genet 102:253–257. doi:10.1007/s004390050688

Hales JB, Brewer JB. 2011. The timing of associative memory formation: frontal lobe and anterior medial temporal lobe activity at associative binding predicts memory. J Neurophysiol 105:1454–1463. doi:10.1152/jn.00902.2010

Hamm V, Héraud C, Bott J-B, Herbeaux K, Strittmatter C, Mathis C, Goutagny R. 2017. Differential contribution of APP metabolites to early cognitive deficits in a TgCRND8 mouse model of Alzheimer‘s disease. Sci Adv 3:e1601068. doi:10.1126/sciadv.1601068

Hampstead BM, Stringer AY, Stilla RF, Amaraneni A, Sathian K. 2011. Where did I put that? Patients with amnestic mild cognitive impairment demonstrate widespread reductions in activity during the encoding of ecologically relevant object-location associations. Neuropsychologia 49:2349–2361. doi:10.1016/j.neuropsychologia.2011.04.008

Hampstead BM, Towler S, Stringer AY, Sathian K. 2018. Continuous measurement of object location memory is sensitive to effects of age and mild cognitive impairment and related to medial temporal lobe volume. Alzheimers Dement (Amst) 10:76–85. doi:10.1016/j.dadm.2017.10.007

Hernandez AR, Reasor JE, Truckenbrod LM, Lubke KN, Johnson SA, Bizon JL, Maurer AP, Burke SN. 2017. Medial prefrontal-perirhinal cortical communication is necessary for flexible response selection. Neurobiology of Learning and Memory 137:36–47. doi:10.1016/j.nlm.2016.10.012

Hirni DI, Kivisaari SL, Krumm S, Monsch AU, Berres M, Oeksuez F, Reinhardt J, Ulmer S, Kressig RW, Stippich C, Taylor KI. 2016. Neuropsychological Markers of Medial Perirhinal and Entorhinal Cortex Functioning are Impaired Twelve Years Preceding Diagnosis of Alzheimer‘s Dementia. JAD 52:573–580. doi:10.3233/JAD-150158

Hyman B, Van Hoesen G, Damasio A, Barnes C. 1984. Alzheimer‘s disease: cell-specific pathology isolates the hippocampal formation. Science 225:1168–1170. doi:10.1126/science.6474172

Jacob P-Y, Casali G, Spieser L, Page H, Overington D, Jeffery K. 2017. An independent, landmark-dominated head-direction signal in dysgranular retrosplenial cortex. Nature Neuroscience 20:173–175. doi:10.1038/nn.4465

Jones DT, Graff-Radford J, Lowe VJ, Wiste HJ, Gunter JL, Senjem ML, Botha H, Kantarci K, Boeve BF, Knopman DS, Petersen RC, Jack CR. 2017. Tau, amyloid, and cascading network failure across the Alzheimer‘s disease spectrum. Cortex, Special Section dedicated to the temporal and parietal lobes 97:143–159. doi:10.1016/j.cortex.2017.09.018

Jones DT, Knopman DS, Gunter JL, Graff-Radford J, Vemuri P, Boeve BF, Petersen RC, Weiner MW, Jack CR Jr, on behalf of the Alzheimer‘s Disease Neuroimaging Initiative. 2016. Cascading network failure across the Alzheimer‘s disease spectrum. Brain 139:547–562. doi:10.1093/brain/awv338

Khan UA, Liu L, Provenzano FA, Berman DE, Profaci CP, Sloan R, Mayeux R, Duff KE, Small SA. 2014. Molecular drivers and cortical spread of lateral entorhinal cortex dysfunction in preclinical Alzheimer‘s disease. Nat Neurosci 17:304–311. doi:10.1038/nn.3606

Kinnavane L, Albasser MM, Aggleton JP. 2015. Advances in the behavioural testing and network imaging of rodent recognition memory. Behavioural Brain Research, SI: Object Recognition Memory in Rats and Mice 285:67–78. doi:10.1016/j.bbr.2014.07.049

Latora V, Marchiori M. 2001. Efficient Behavior of Small-World Networks. Phys Rev Lett 87:198701. doi:10.1103/PhysRevLett.87.198701

Lein ES, Hawrylycz MJ, Ao N, Ayres M, Bensinger A, Bernard A, Boe AF, Boguski MS, Brockway KS, Byrnes EJ, Chen Lin, Chen Li, Chen T-M, Chi Chin M, Chong J, Crook BE, Czaplinska A, Dang CN, Datta S, Dee NR, Desaki AL, Desta T, Diep E, Dolbeare TA, Donelan MJ, Dong H-W, Dougherty JG, Duncan BJ, Ebbert AJ, Eichele G, Estin LK, Faber C, Facer BA, Fields R, Fischer SR, Fliss TP, Frensley C, Gates SN, Glattfelder KJ, Halverson KR, Hart MR, Hohmann JG, Howell MP, Jeung DP, Johnson RA, Karr PT, Kawal R, Kidney JM, Knapik RH, Kuan CL, Lake JH, Laramee AR, Larsen KD, Lau C, Lemon TA, Liang AJ, Liu Y, Luong LT, Michaels J, Morgan JJ, Morgan RJ, Mortrud MT, Mosqueda NF, Ng LL, Ng R, Orta GJ, Overly CC, Pak TH, Parry SE, Pathak SD, Pearson OC, Puchalski RB, Riley ZL, Rockett HR, Rowland SA, Royall JJ, Ruiz MJ, Sarno NR, Schaffnit K, Shapovalova NV, Sivisay T, Slaughterbeck CR, Smith SC, Smith KA, Smith BI, Sodt AJ, Stewart NN, Stumpf K-R, Sunkin SM, Sutram M, Tam A, Teemer CD, Thaller C, Thompson CL, Varnam LR, Visel A, Whitlock RM, Wohnoutka PE, Wolkey CK, Wong VY, Wood M, Yaylaoglu MB, Young RC, Youngstrom BL, Feng Yuan X, Zhang B, Zwingman TA, Jones AR. 2007. Genome-wide atlas of gene expression in the adult mouse brain. Nature 445:168–176. doi:10.1038/nature05453

Liang J, Li Y, Liu H, Zhang S, Wang M, Chu Y, Ye J, Xi Q, Zhao X. 2020. Increased intrinsic default-mode network activity as a compensatory mechanism in aMCI: a resting-state functional connectivity MRI study. Aging (Albany NY) 12:5907–5919. doi:10.18632/aging.102986

Lin S-Y, Lin C-P, Hsieh T-J, Lin C-F, Chen S-H, Chao Y-P, Chen Y-S, Hsu C-C, Kuo L-W. 2019. Multiparametric graph theoretical analysis reveals altered structural and functional network topology in Alzheimer‘s disease. NeuroImage: Clinical 22:101680. doi:10.1016/j.nicl.2019.101680

Liu Z, Zhang Y, Yan H, Bai L, Dai R, Wei W, Zhong C, Xue T, Wang H, Feng Y, You Y, Zhang X, Tian J. 2012. Altered topological patterns of brain networks in mild cognitive impairment and Alzheimer‘s disease: A resting-state fMRI study. Psychiatry Research: Neuroimaging 202:118–125. doi:10.1016/j.pscychresns.2012.03.002

Lu H, Zou Q, Gu H, Raichle ME, Stein EA, Yang Y. 2012. Rat brains also have a default mode network. PNAS 109:3979–3984. doi:10.1073/pnas.1200506109

Maass A, Berron D, Harrison TM, Adams JN, La Joie R, Baker S, Mellinger T, Bell RK, Swinnerton K, Inglis B, Rabinovici GD, Düzel E, Jagust WJ. 2019. Alzheimer‘s pathology targets distinct memory networks in the ageing brain. Brain 142:2492–2509. doi:10.1093/brain/awz154

McAvoy K, Besnard A, Sahay A. 2015. Adult hippocampal neurogenesis and pattern separation in DG: a role for feedback inhibition in modulating sparseness to govern population-based coding. Front Syst Neurosci 9. doi:10.3389/fnsys.2015.00120

Miller AMP, Vedder LC, Law LM, Smith DM. 2014. Cues, context, and long-term memory: the role of the retrosplenial cortex in spatial cognition. Front Hum Neurosci 8. doi:10.3389/fnhum.2014.00586

Mitchell AS, Czajkowski R, Zhang N, Jeffery K, Nelson AJD. 2018. Retrosplenial cortex and its role in spatial cognition. Brain Neurosci Adv 2. doi:10.1177/2398212818757098

Morys J, Bobinski M, Wegiel J, Wisniewski HM, Narkiewicz O. 1996. Alzheimer’s disease severely affects areas of the claustrum connected with the entorhinal cortex. J Hirnforsch 37:173–180.

Myers N, Pasquini L, Göttler J, Grimmer T, Koch K, Ortner M, Neitzel J, Mühlau M, Förster S, Kurz A, Förstl H, Zimmer C, Wohlschläger AM, Riedl V, Drzezga A, Sorg C. 2014. Within-patient correspondence of amyloid-β and intrinsic network connectivity in Alzheimer‘s disease. Brain 137:2052–2064. doi:10.1093/brain/awu103

Naveh-Benjamin M. 2000. Adult age differences in memory performance: Tests of an associative deficit hypothesis. Journal of Experimental Psychology: Learning, Memory, and Cognition 26:1170–1187. doi:10.1037/0278-7393.26.5.1170

Nuriel T, Angulo SL, Khan U, Ashok A, Chen Q, Figueroa HY, Emrani S, Liu L, Herman M, Barrett G, Savage V, Buitrago L, Cepeda-Prado E, Fung C, Goldberg E, Gross SS, Hussaini SA, Moreno H, Small SA, Duff KE. 2017. Neuronal hyperactivity due to loss of inhibitory tone in APOE4 mice lacking Alzheimer‘s disease-like pathology. Nat Commun 8:1464. doi:10.1038/s41467-017-01444-0

Ogomori K, Kitamoto T, Tateishi J, Sato Y, Suetsugu M, Abe M. 1989. Beta-protein amyloid is widely distributed in the central nervous system of patients with Alzheimer‘s disease. Am J Pathol 134:243–251.

Palmqvist S, Schöll M, Strandberg O, Mattsson N, Stomrud E, Zetterberg H, Blennow K, Landau S, Jagust W, Hansson O. 2017. Earliest accumulation of β-amyloid occurs within the default-mode network and concurrently affects brain connectivity. Nature Communications 8:1214. doi:10.1038/s41467-017-01150-x

Parron C, Save E. 2004. Comparison of the effects of entorhinal and retrosplenial cortical lesions on habituation, reaction to spatial and non-spatial changes during object exploration in the rat. Neurobiology of Learning and Memory 82:1–11. doi:10.1016/j.nlm.2004.03.004

Pereira JB, Mijalkov M, Kakaei E, Mecocci P, Vellas B, Tsolaki M, Kłoszewska I, Soininen H, Spenger C, Lovestone S, Simmons A, Wahlund L-O, Volpe G, Westman E. 2016. Disrupted Network Topology in Patients with Stable and Progressive Mild Cognitive Impairment and Alzheimer‘s Disease. Cereb Cortex 26:3476–3493. doi:10.1093/cercor/bhw128

Pereira JB, Ossenkoppele R, Palmqvist S, Strandberg TO, Smith R, Westman E, Hansson O. 2019. Amyloid and tau accumulate across distinct spatial networks and are differentially associated with brain connectivity. eLife 8:e50830. doi:10.7554/eLife.50830

Perusini JN, Cajigas SA, Cohensedgh O, Lim SC, Pavlova IP, Donaldson ZR, Denny CA. 2017. Optogenetic stimulation of dentate gyrus engrams restores memory in Alzheimer‘s disease mice. Hippocampus 27:1110–1122. doi:10.1002/hipo.22756

Petersen RC. 2011. Mild Cognitive Impairment. New England Journal of Medicine 364:2227–2234. doi:10.1056/NEJMcp0910237

Petrache AL, Rajulawalla A, Shi A, Wetzel A, Saito T, Saido TC, Harvey K, Ali AB. 2019. Aberrant Excitatory–Inhibitory Synaptic Mechanisms in Entorhinal Cortex Microcircuits During the Pathogenesis of Alzheimer‘s Disease. Cerebral Cortex 29:1834–1850. doi:10.1093/cercor/bhz016

Pusil S, López ME, Cuesta P, Bruña R, Pereda E, Maestú F. 2019. Hypersynchronization in mild cognitive impairment: the ‘X’ model. Brain 142:3936–3950. doi:10.1093/brain/awz320

Qin Y-Y, Li M-W, Zhang S, Zhang Y, Zhao L-Y, Lei H, Oishi K, Zhu W-Z. 2013. In vivo quantitative whole-brain diffusion tensor imaging analysis of APP/PS1 transgenic mice using voxel-based and atlas-based methods. Neuroradiology 55:1027–1038. doi:10.1007/s00234-013-1195-0

Reagh ZM, Noche JA, Tustison NJ, Delisle D, Murray EA, Yassa MA. 2018. Functional Imbalance of Anterolateral Entorhinal Cortex and Hippocampal Dentate/CA3 Underlies Age-Related Object Pattern Separation Deficits. Neuron 97:1187–1198.e4. doi:10.1016/j.neuron.2018.01.039

Reagh ZM, Yassa MA. 2014. Object and spatial mnemonic interference differentially engage lateral and medial entorhinal cortex in humans. Proceedings of the National Academy of Sciences 111:E4264–E4273. doi:10.1073/pnas.1411250111

Rentz DM, Amariglio RE, Becker JA, Frey M, Olson LE, Frishe K, Carmasin J, Maye JE, Johnson KA, Sperling RA. 2011. Face-name associative memory performance is related to amyloid burden in normal elderly. Neuropsychologia 49:2776–2783. doi:10.1016/j.neuropsychologia.2011.06.006

Ritchey M, Cooper RA. 2020. Deconstructing the Posterior Medial Episodic Network. Trends in Cognitive Sciences 24:451–465. doi:10.1016/j.tics.2020.03.006

Rodriguez GA, Barrett GM, Duff KE, Hussaini SA. 2020. Chemogenetic attenuation of neuronal activity in the entorhinal cortex reduces Aβ and tau pathology in the hippocampus. PLOS Biology 18:e3000851. doi:10.1371/journal.pbio.3000851

Roy DS, Arons A, Mitchell TI, Pignatelli M, Ryan TJ, Tonegawa S. 2016. Memory retrieval by activating engram cells in mouse models of early Alzheimer‘s disease. Nature 531:508–512. doi:10.1038/nature17172

Rubinov M, Sporns O. 2010. Complex network measures of brain connectivity: Uses and interpretations. NeuroImage, Computational Models of the Brain 52:1059–1069. doi:10.1016/j.neuroimage.2009.10.003

Saito T, Matsuba Y, Mihira N, Takano J, Nilsson P, Itohara S, Iwata N, Saido TC. 2014. Single App knock-in mouse models of Alzheimer‘s disease. Nat Neurosci 17:661–663. doi:10.1038/nn.3697

Saito T, Mihira N, Matsuba Y, Sasaguri H, Hashimoto S, Narasimhan S, Zhang B, Murayama S, Higuchi M, Lee VMY, Trojanowski JQ, Saido TC. 2019. Humanization of the entire murine *Mapt* gene provides a murine model of pathological human tau propagation. J Biol Chem jbc.RA119.009487. doi:10.1074/jbc.RA119.009487

Sanabria A, Alegret M, Rodriguez-Gomez O, Valero S, Sotolongo-Grau O, Monté-Rubio G, Abdelnour C, Espinosa A, Ortega G, Perez-Cordon A, Gailhajanet A, Hernandez I, Rosende-Roca M, Vargas L, Mauleon A, Sanchez D, Martin E, Rentz DM, Lomeña F, Ruiz A, Tarraga L, Boada M. 2018. The Spanish version of Face-Name Associative Memory Exam (S-FNAME) performance is related to amyloid burden in Subjective Cognitive Decline. Scientific Reports 8:3828. doi:10.1038/s41598-018-21644-y

Sasaguri H, Nilsson P, Hashimoto S, Nagata K, Saito T, Strooper BD, Hardy J, Vassar R, Winblad B, Saido TC. 2017. APP mouse models for Alzheimer‘s disease preclinical studies. The EMBO Journal 36:2473–2487. doi:10.15252/embj.201797397

Scearce-Levie K, Sanchez PE, Lewcock JW. 2020. Leveraging preclinical models for the development of Alzheimer disease therapeutics. Nature Reviews Drug Discovery 19:447–462. doi:10.1038/s41573-020-0065-9

Schultz AP, Chhatwal JP, Hedden T, Mormino EC, Hanseeuw BJ, Sepulcre J, Huijbers W, LaPoint M, Buckley RF, Johnson KA, Sperling RA. 2017. Phases of hyper and hypo connectivity in the Default Mode and Salience networks track with amyloid and Tau in clinically normal individuals. J Neurosci. doi:10.1523/JNEUROSCI.3263-16.2017

Selkoe DJ. 2012. Preventing Alzheimer‘s Disease. Science 337:1488–1492. doi:10.1126/science.1228541

Sepulcre J, Sabuncu MR, Li Q, El Fakhri G, Sperling R, Johnson KA. 2017. Tau and amyloid β proteins distinctively associate to functional network changes in the aging brain. Alzheimer’s & Dementia 13:1261–1269. doi:10.1016/j.jalz.2017.02.011

Shah D, Latif-Hernandez A, Strooper B, Saito T, Saido T, Verhoye M, D‘Hooge R, Linden A. 2018. Spatial reversal learning defect coincides with hypersynchronous telencephalic BOLD functional connectivity in APP NL-F/NL-F knock-in mice. Scientific Reports 8:6264. doi:10.1038/s41598-018-24657-9

Shah D, Praet J, Latif Hernandez A, Höfling C, Anckaerts C, Bard F, Morawski M, Detrez JR, Prinsen E, Villa A, De Vos WH, Maggi A, D‘Hooge R, Balschun D, Rossner S, Verhoye M, Van der Linden A. 2016. Early pathologic amyloid induces hypersynchrony of BOLD resting-state networks in transgenic mice and provides an early therapeutic window before amyloid plaque deposition. Alzheimer’s & Dementia 12:964–976. doi:10.1016/j.jalz.2016.03.010

Sinha N, Berg CN, Tustison NJ, Shaw A, Hill D, Yassa MA, Gluck MA. 2018. APOE ε4 Status in Healthy Older African Americans is Associated with Deficits in Pattern Separation and Hippocampal Hyperactivation. Neurobiol Aging 69:221–229. doi:10.1016/j.neurobiolaging.2018.05.023

Stafford JM, Jarrett BR, Miranda-Dominguez O, Mills BD, Cain N, Mihalas S, Lahvis GP, Lattal KM, Mitchell SH, David SV, Fryer JD, Nigg JT, Fair DA. 2014. Large-scale topology and the default mode network in the mouse connectome. PNAS 111:18745–18750. doi:10.1073/pnas.1404346111

Sugar J, Witter MP, van Strien N, Cappaert N. 2011. The Retrosplenial Cortex: Intrinsic Connectivity and Connections with the (Para)Hippocampal Region in the Rat. An Interactive Connectome. Front Neuroinform 5. doi:10.3389/fninf.2011.00007

Tanimizu T, Kenney JW, Okano E, Kadoma K, Frankland PW, Kida S. 2017. Functional Connectivity of Multiple Brain Regions Required for the Consolidation of Social Recognition Memory. J Neurosci 37:4103–4116. doi:10.1523/JNEUROSCI.3451-16.2017

Tischmeyer W, Grimm R. 1999. Activation of immediate early genes and memory formation. Cell Mol Life Sci 55:564–574. doi:10.1007/s000180050315

Vann SD, Aggleton JP. 2005. Selective dysgranular retrosplenial cortex lesions in rats disrupt allocentric performance of the radial-arm maze task. Behav Neurosci 119:1682–1686. doi:10.1037/0735-7044.119.6.1682

Vann SD, Aggleton JP, Maguire EA. 2009. What does the retrosplenial cortex do? Nature Reviews Neuroscience 10:792–802. doi:10.1038/nrn2733

Vetere G, Kenney JW, Tran LM, Xia F, Steadman PE, Parkinson J, Josselyn SA, Frankland PW. 2017. Chemogenetic Interrogation of a Brain-wide Fear Memory Network in Mice. Neuron 94:363–374.e4. doi:10.1016/j.neuron.2017.03.037

Vogt BA, Paxinos G. 2014. Cytoarchitecture of mouse and rat cingulate cortex with human homologies. Brain Struct Funct 219:185–192. doi:10.1007/s00429-012-0493-3

Wang J, Zuo X, Dai Z, Xia M, Zhao Z, Zhao X, Jia J, Han Y, He Y. 2013a. Disrupted Functional Brain Connectome in Individuals at Risk for Alzheimer‘s Disease. Biological Psychiatry, Disturbances in the Connectome and Risk for Alzheimer‘s Disease 73:472–481. doi:10.1016/j.biopsych.2012.03.026

Wang JX, Rogers LM, Gross EZ, Ryals AJ, Dokucu ME, Brandstatt KL, Hermiller MS, Voss JL. 2014. Targeted enhancement of cortical-hippocampal brain networks and associative memory. Science 345:1054–1057. doi:10.1126/science.1252900

Wang L, Brier MR, Snyder AZ, Thomas JB, Fagan AM, Xiong C, Benzinger TL, Holtzman DM, Morris JC, Ances BM. 2013b. Cerebrospinal Fluid Aβ42, Phosphorylated Tau _181_, and Resting-State Functional Connectivity. JAMA Neurol. doi:10.1001/jamaneurol.2013.3253

Wang Q, Ng L, Harris JA, Feng D, Li Y, Royall JJ, Oh SW, Bernard A, Sunkin SM, Koch C, Zeng H. 2017. Organization of the connections between claustrum and cortex in the mouse. J Comp Neurol 525:1317–1346. doi:10.1002/cne.24047

Wang Y, Risacher SL, West JD, McDonald BC, Magee TR, Farlow MR, Gao S, O‘Neill DP, Saykin AJ. 2013c. Altered default mode network connectivity in older adults with cognitive complaints and amnestic mild cognitive impairment. J Alzheimers Dis 35:751–760. doi:10.3233/JAD-130080

Warren KN, Hermiller MS, Nilakantan AS, Voss JL. 2019. Stimulating the hippocampal posterior-medial network enhances task-dependent connectivity and memory. eLife 8:e49458. doi:10.7554/eLife.49458

Weiler M, Stieger KC, Long JM, Rapp PR. 2020. Transcranial Magnetic Stimulation in Alzheimer‘s Disease: Are We Ready? eNeuro 7. doi:10.1523/ENEURO.0235-19.2019

Wheeler AL, Teixeira CM, Wang AH, Xiong X, Kovacevic N, Lerch JP, McIntosh AR, Parkinson J, Frankland PW. 2013. Identification of a Functional Connectome for Long-Term Fear Memory in Mice. PLOS Computational Biology 9:e1002853. doi:10.1371/journal.pcbi.1002853

Wilson DIG, Langston RF, Schlesiger MI, Wagner M, Watanabe S, Ainge JA. 2013a. Lateral entorhinal cortex is critical for novel object-context recognition. Hippocampus 23:352–366. doi:10.1002/hipo.22095

Wilson DIG, Watanabe S, Milner H, Ainge JA. 2013b. Lateral entorhinal cortex is necessary for associative but not nonassociative recognition memory. Hippocampus 23:1280–1290. doi:10.1002/hipo.22165

Wright DB, London K, Field AP. 2011. Using Bootstrap Estimation and the Plug-in Principle for Clinical Psychology Data. Journal of Experimental Psychopathology 2:252–270. doi:10.5127/jep.013611

Xu W, Fitzgerald S, Nixon RA, Levy E, Wilson DA. 2015. Early hyperactivity in lateral entorhinal cortex is associated with elevated levels of AβPP metabolites in the Tg2576 mouse model of Alzheimer‘s disease. Exp Neurol 264:82–91. doi:10.1016/j.expneurol.2014.12.008

Yassa MA, Stark SM, Bakker A, Albert MS, Gallagher M, Stark CEL. 2010. High-resolution structural and functional MRI of hippocampal CA3 and dentate gyrus in patients with amnestic Mild Cognitive Impairment. NeuroImage 51:1242–1252. doi:10.1016/j.neuroimage.2010.03.040

Yeung L-K, Olsen RK, Hong B, Mihajlovic V, D‘Angelo MC, Kacollja A, Ryan JD, Barense MD. 2018. Object-in-Place Memory Predicted by Anterolateral Entorhinal Cortex and Parahippocampal Cortex Volume in Older Adults. bioRxiv 409607. doi:10.1101/409607

Zott B, Simon MM, Hong W, Unger F, Chen-Engerer H-J, Frosch MP, Sakmann B, Walsh DM, Konnerth A. 2019. A vicious cycle of β amyloid–dependent neuronal hyperactivation. Science 365:559–565. doi:10.1126/science.aay0198

